# Comparing cortex-wide gene expression patterns between species in a common reference frame

**DOI:** 10.1101/2021.07.28.454203

**Authors:** Mackenzie Englund, Sebastian S. James, Riley Bottom, Kelly J. Huffman, Stuart P. Wilson, Leah A. Krubitzer

## Abstract

Advances in sequencing techniques have made comparative studies of gene expression a current focus for understanding evolutionary and developmental processes. However, insights into the spatial expression of genes have been limited by a lack of robust methodology. We therefore developed a set of algorithms for quantifying and comparing tissue-wide spatial patterns of gene expression within and across species. Here we apply these algorithms to compare cortex-wide expression of *Id2* and *RZRβ* mRNA in early postnatal mice and voles. We show that neocortical patterns of *Id2* expression are moderately conserved between species, but that the degree of conservation varies by cortical layer and area. By comparison, patterns of *RZRβ* expression are highly conserved in somatosensory areas, and more variable between species in visual and auditory areas. We consider if these differences reflect independent evolution in the 35 million years since the last common ancestor.

---

Almost everything we know about the human brain comes from comparative studies of other animals: from genes involved in cortical development to system-level networks that generate complex behaviours. Comparative studies of living species provide a robust means by which to understand unknown forms, like humans, and even extinct forms like our early mammalian ancestors. Importantly, these types of studies are critical for identifying features of brain organization that are conserved between species and those that may have been derived in different lineages. They also allow us to determine how developmental programs and timing schedules may vary across species, and better understand how phenotypic diversity can be generated over shorter and longer timescales. Finally, by making valid comparisons across species, we can begin to understand how complexity emerges in different nervous systems, the rules of brain construction, and the constraints imposed on developing and evolving nervous systems.

Despite the importance of comparative studies in biology, most comparisons of anatomically reconstructed data are subjective, and most gene sequencing studies neglect the actual spatial patterns of gene expression across a structure, focusing instead on cell-type expression (*1–3*). Moreover, many current methods for making comparisons fail to capture the three-dimensional nature of the brain, which is composed of asymmetrical structures that can vary markedly in relative shape, size, and location across species and between developmental time-points. Despite the 3D nature of the brain, most studies collapse data into two dimensions driven largely by the plane of section at which the brain is cut.

As such, neurobiologists are faced with two challenges. First, attempting to understand 3D structures by analyzing 2D images is inherently problematic because the loss of spatial information is unavoidable, especially in curved structures (*4*). 2D analysis often involves pre-specifying regions of interest (ROI’s) to quantify the presence of labelled cells or mRNA expression after in-situ hybridization (ISH), narrowing the focus and potentially missing overall differences. A second challenge, which arises when making comparisons between structures in different species and/or at different developmental time-points, or between different experimental conditions, is determining the extent to which 2D spatial patterning might be invariant to basic transformations in the size and shape of the 3D structure. To this end, it is important for comparisons to be made with respect to a common anatomical reference frame.

In the current study, we overcame these challenges by developing a set of algorithms for brain slice registration in 3D, and for incorporating ISH data into a common reference frame to enable point-by-point comparisons between species or experimental conditions. These tools, which we refer to collectively as *Stalefish*, *The Spatial Analysis of Fluorescent (and non-fluorescent) In-Situ Hybridization*, allowed for the laminar and spatial patterns of expression of genes involved in cortical development to be quantified and compared in two species of age matched rodents.

Our analysis of *Id2* and *RZRβ* cortical expression patterns in early postnatal mouse and vole brains reveals both a strong layer-specific conservation of the patterning of these genes, as well as area-specific differences that shed new light on the ontogeny and phylogeny of neocortical arealization.

## Reconstructing whole-brain patterns of gene expression from processed tissue

In mice (*Mus musculus)* and prairie voles (*Microtus ochrogaster*), direct layer- and area-specific comparisons were made between the cortex-wide expression patterns of two genes important for cortical development: *Id2* (Inhibitor of DNA-binding 2), and *RZRβ* (RAR-related orphan receptor beta) (*5–9*). Using a novel algorithm for slice registration developed as part of the *Stalefish* methodology, we could for the first time visualize and quantify cortex-wide layer-specific patterns of gene expression in 3D, and determine the extent to which these patterns are conserved across species or derived in a particular lineage. To illustrate how layer-specific cortical in-situ hyridization (ISH) data can be reconstructed from coronal sections, the *Stalefish* curve drawing tool was first used to process an entire hemisphere of a postnatal day zero (P0) vole brain (Fig. 1A) hybridized for *Id2* (Fig. 1, B-E). The *Stalefish* 3D viewer tool was then used to show the 3D-reconstructed expression pattern in the superficial layers (Fig. 1, F-G). Finally, the *Stalefish* digital flattening tool was used to project the data into a new 2D plane for subsequent analysis. In this plane, the piriform cortex became clearly demarcated as an area of high expression, and the shapes and positions of several other areas of high and low expression in the neonatal vole cortex were revealed to correspond well with descriptions of cortical field boundaries later observed in adult voles (*10*) (Fig. 1H), confirming the proposed role of this gene as an ‘area marker’ for putative neocortical fields. The *Stalefish* methodology was further validated by comparing reconstructions of our mouse data to reconstructions that we made using similar data obtained from the Allen Mouse Brain Atlas (*11, 12*) (fig. S1).

**Fig. 1.**
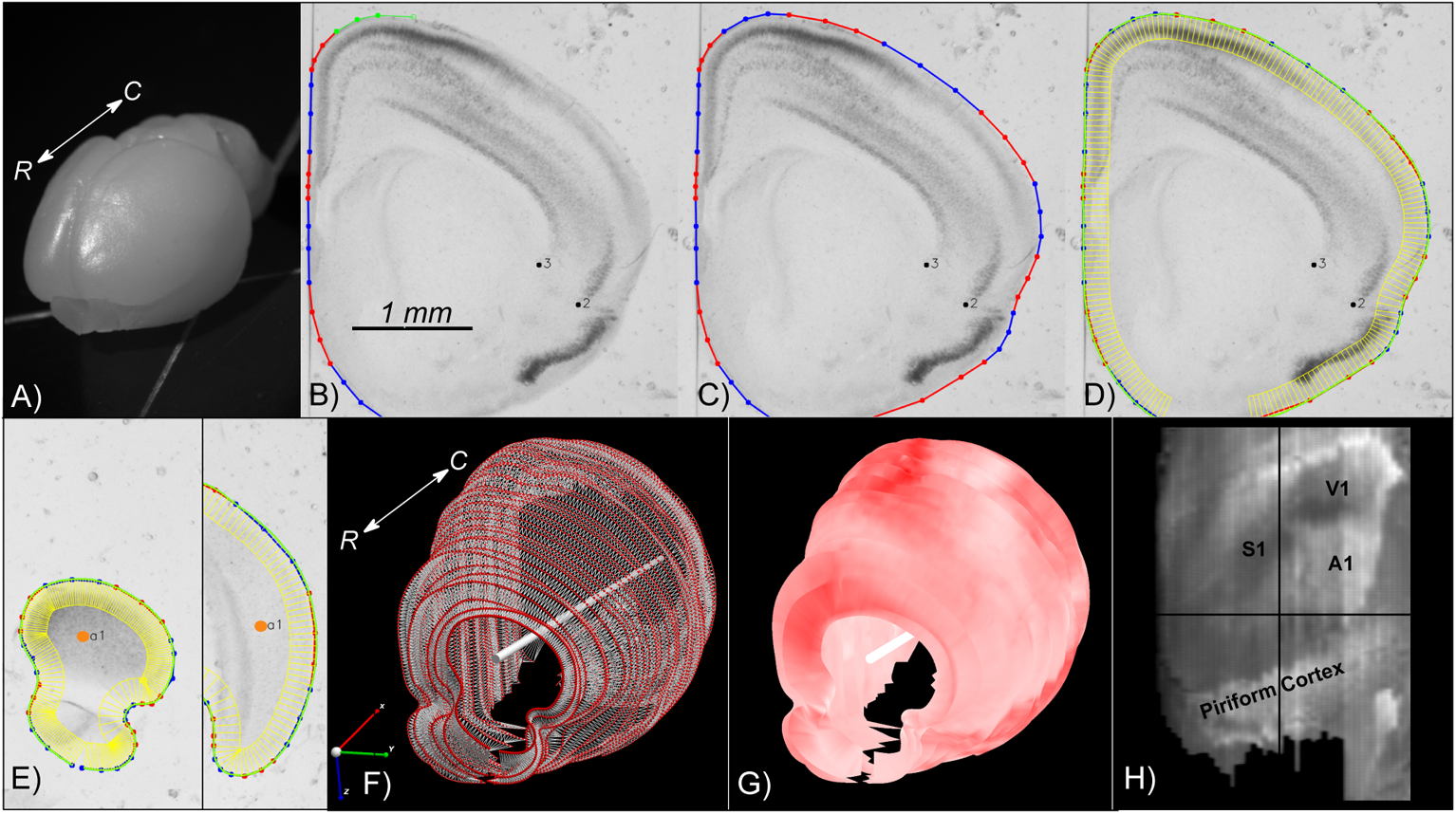
Stalefish Workflow. (A) Rostral view of an unfixed vole brain. The rostral-caudal axis is shown (white arrow). (B-E) Screenshots from *Stalefish* of coronal sections of *Id2*-hybridized vole tissue showing the curve drawing method. Dark regions indicate a higher incidence of *Id2* mRNA. (B) Marked locations around the perimeter of the brain. The perimeter points are collected into small sets of 4 or 5 points at a time. The green points are the newest set of perimeter locations and will become the next ‘blue’ set (the red and blue colors are simply a guide for the user). The number of points in each section determines the order of the Bezier curve which will be fitted to that section. The black dots numbered 2 and 3 show landmark placement location for alignment. (C) Once the perimeter points have been laid out, a piecewise fit is found for the points by modifying the individual Bezier curves to ensure that the curve gradient is continuous at the joints. (D) The green line shows the final fit. Evenly spaced normal vectors extended down from the fit line give sampling boxes in yellow. For each sampling box, the user can access the mean luminance, along with all of the per-pixel values and the position within the sampling box. (E) An axis mark (orange dot, drawn larger for illustrative purpose) on the first and last slice define a user-defined ‘brain axis’ used for alignment. (F) *Stalefish* output using sfview. Curve points are shown as red spheres. By connecting the spheres to make a mesh, a surface is generated. The white bar shows the user-defined brain axis, derived from the two axis marks placed on the brain slices. The rostral-caudal axis is shown (white arrow). (G) The mean luminance of the sampling boxes can then be indicated on the smoothed surface to give a 3D reconstruction of the gene expression map. Here, we used a monochrome colormap for which full-saturation red corresponds to the maximum *Id2* expression signal. (H) Digitally-flattened and reference-frame transformed 3D surface map (from B-G) using sfview. Abbreviations: rostral (R), caudal (C), millimeter (mm), primary somatosensory cortex (S1), primary visual cortex (V1), primary auditory cortex (A1).

## Conservation of cortex-wide gene expression across species

*Id2* is a key transcription factor in neurodevelopment that regulates cellular differentiation and neurite outgrowth (*13–16*). We quantified the degree of conservation of cortex-wide patterns of *Id2* expression between prairie voles and mice and found that the laminar and areal patterns of *Id2* expression are highly conserved across species. We first hybridized neonatal vole (n = 7) and mouse (n = 3) brain tissue for *Id2* in animals matched for developmental stage. *Id2* was expressed in all cortical layers except layer 4 in both species, similar to findings from previous studies (*17*). *Stalefish* was used to fit curves on each *Id2*-hybridized section of the series at depths from the neocortical surface that correspond to layer 2/3, layer 5, and layer 6, and to digitally unwrap and flatten the 3D reconstructed expression patterns. Using the positions of external anatomical landmark locations commonly identifiable in all brains, a linear transformation was computed for mapping the flattened reconstruction of each brain to the coordinate system of an arbitrary reference brain from the comparison set, allowing cortex-wide expression patterns from multiple brains to be compared point-by-point in a common reference frame (See *Methods*).

Following these transformations, we quantified the similarity of cortex-wide patterns of *Id2* expression by applying Pearson correlation analyses to the point-by-point matched expression levels between maps, excluding any points not present in every pattern submitted for a given set of comparisons. Correlation coefficients were calculated for each pair of expression patterns and comparisons were made with respect to cortical layer, and within and across species (Fig. 2, A and B). To our knowledge, this is the first time that cortex-wide patterns have ever been quantitatively compared across layers and species. Within-species comparisons demonstrated that layer 2/3 expression maps were highly correlated (Vole: r_avg_(6) = .523 ± .105, Mouse: r_avg_(3) = .854 ± .031), as were layer 5 and layer 6 maps (Layer 5: Vole: r_avg_(6) = .605 ± .077, Mouse: r_avg_(3) = .873 ± .015) (Layer 6: Vole: r_avg_(6) = .748 ± .047, Mouse: r_avg_(3) = .644 ± .041). Examining expression maps across layers within a species revealed that layer 2/3 expression was weakly correlated with layer 5 (Vole: r_avg_(16) = .159 ± .141, Mouse: r_avg_(9) = .318 ± .078), and layer 5 expression was weakly correlated with layer 6 expression (Vole: r_avg_(16) = .169 ± .078, Mouse: r_avg_(9) = .303 ± .075). Conversely, as expected by the lack of expression in layer 2/3 of primary somatosensory (S1) and primary visual (V1) cortex, and high expression in layer 6 of these areas, layer 2/3 expression was negatively correlated with layer 6 expression in both mice and voles (Vole: r_avg_(16) = −.208 ± .149, Mouse: r_avg_(9) = −.241 ± .073). As depicted in Figure 2A and 2B, the directions of these layer-specific correlations were remarkably well conserved across species. However, when comparing the degree of within-layer correlation across species, mice exhibited significantly higher correlations on 5 out of the 6 comparisons (p_Layer 2/3_ = .014, p_Layer 5_ = .014, p_Layer 6_ = .047, p_Layer 2/3 vs 5_ = .007, p_Layer 2/3 vs 6_ = .466, p_Layer 5 vs 6_ = .001), suggesting that spatial patterns of expression of *Id2* amongst neonatal mice varied less than amongst neonatal voles. Between species, layer 2/3 *Id2* expression in mice (n = 3) and voles (n = 4) was positively correlated (r_avg_(12) = .523 ± .109) (Fig. 2C), as were layer 5 and layer 6 expression maps (Layer 5: r_avg_(12) = .482 ± .112, Layer 6: r_avg_(12) = .234 ± .084). Taken together, the similarities in direction and magnitude of the above correlational data provide the first ever direct and quantitative evidence that mice and voles have conserved laminar expression profiles of *Id2* during early postnatal brain development. These findings add support for our hypothesis that the spatial pattern of *Id2* expression across the entire neocortex is conserved on a by-layer basis.

**Fig. 2.**
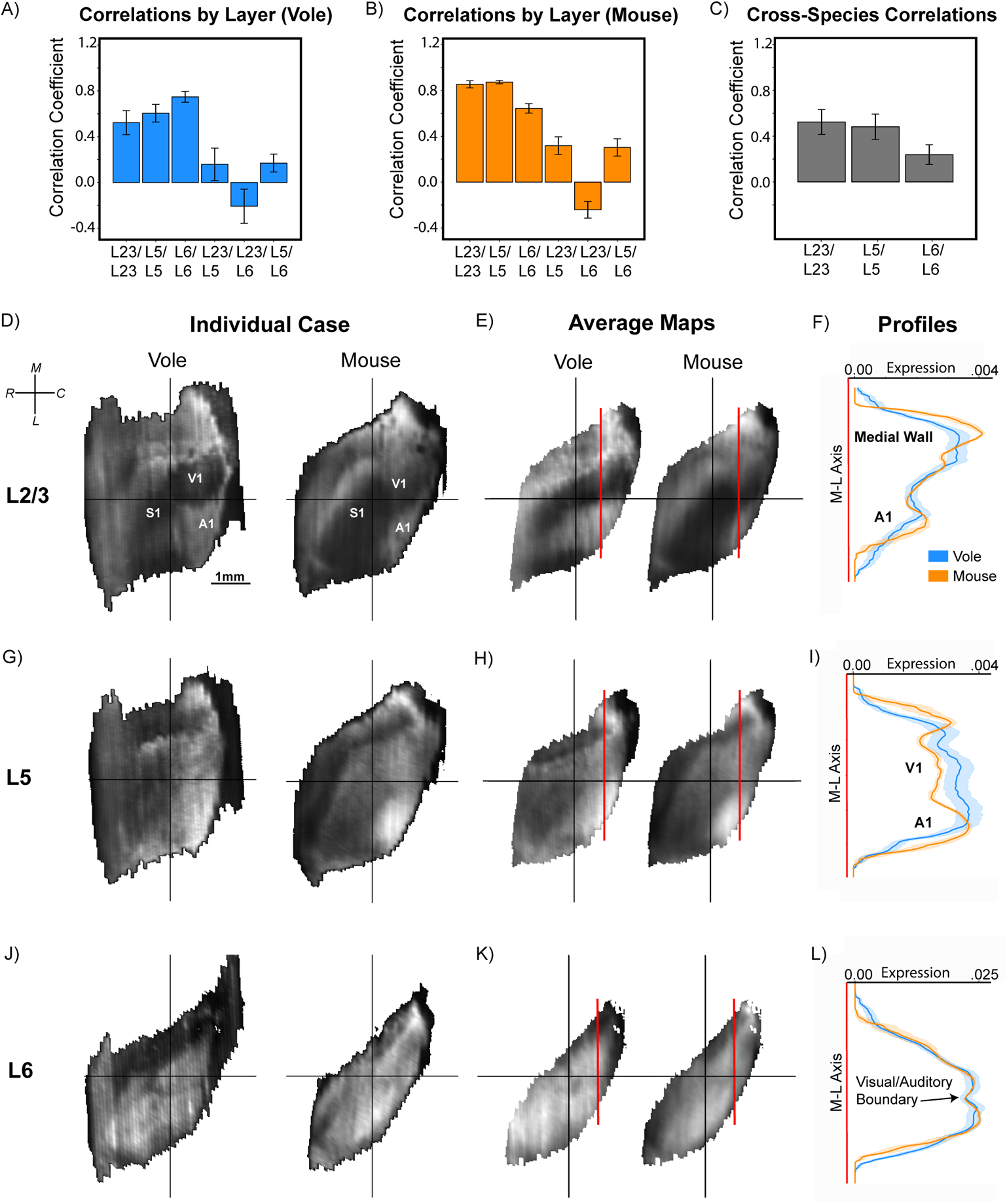
The spatial pattern of *Id2* expression is conserved on a layer-by-layer and area basis. (A & B) Bar graphs of average correlation coefficients between individuals within-species shown as mean with standard deviation across cortical layers. There is little within-species variation of *Id2* expression by layer, and both voles (n=6) and mice (n=3) show similarities in degree and direction of correlations across layers. (C) Bar graph of average correlation coefficient of *Id2* expression maps between species by cortical layer. (D,G,J) Digitally flattened maps (individual example cases) of Id2 expression in different layers reconstructed from coronal sections in a vole (left) and mouse (right) by *Stalefish*. Cases are displayed post-transform and aligned to a common axis. The x and y axes denote spatial coordinates, while expression level is represented as pixel-value at a given coordinate (black = lowest, white = highest). (E, H, K) Average maps of each species by layer. Average maps are displayed as only the locations which overlapped between species post-transform (rostral left, medial up). Vertical red lines show the location at which medial-to-lateral expression profiles were sampled and analyzed (line graphs on the far right). (F, I, L) Line graphs show the 200µM average medial-to-lateral expression profiles between voles (blue) and mice (orange) for a given layer at a specific rostral-caudal depth (red lines in F/I/L). Data is presented as mean with bootstrapped 95% confidence intervals. The x-axis denotes signal level (after mean-centering) and the y-axis denotes the medial-lateral axis from the subiculum to the rhinal fissure. Abbreviations: Layer 2/3 (L23), Layer 5 (L5), Layer 6 (L6), rostral (R), caudal (C), medial (M), lateral (L), millimeter (mm), primary somatosensory cortex (S1), primary visual cortex (V1), primary auditory cortex (A1)

Following our analysis of *Id2* expression by layer, we sought to determine if expression of Id2 in specific regions of the neocortex was conserved across species (Fig. 2, D-L). While previous work has suggested that high *Id2* expression overlaps roughly with sensory areas (*16,18–20*), our data show that extremely precise regions of high and low expression of *Id2* correspond with the shapes of putative cortical areas in each layer, and how this correspondence varies between species. Specifically, in both species there was a marked absence of *Id2* expression in layer 2/3 in the putative primary visual (V1) and primary somatosensory cortex (S1), while *Id2* expression was highly expressed in layer 2/3 of the putative primary auditory cortex (A1) (Figure 2D/E/F). Layer 5 *Id2* expression in both mice and voles was strongest in A1 and in the caudal portion of the medial wall (Fig. 2, G, H, I). In layer 6, *Id2* was highly expressed in all of the putative primary sensory areas (S1, V1, A1), in both voles and mice, although putative cortical areas were more clearly defined in voles (Fig. 2, J, K, L).

Next, we observed that expression maps appeared more similar in the caudal compared to rostral half of the neocortex. To quantify this observation, and to demonstrate the usefulness of the transformed data for comparing expression levels in specific regions of interest, we created 200µM thick expression profiles of the different layers in the caudal portion of the neocortex (Fig. 2, F, I, J). Profiles spanned medial to lateral, encompassing portions of the medial wall, visual, and auditory cortex (red line in Fig. 2 E, H, K). The average expression profiles in voles (blue trace) and mice (orange trace) were highly correlated for layer 2/3 (r(10) = .860), with both species exhibiting high expression at the medial wall, a substantial decrease in expression in putative V1, and a rise in expression in putative A1 (Fig. 2F). The layer 5 profiles showed three clear peaks of expression in both species, corresponding to the medial wall, putative V1, and putative A1, and were also strongly correlated (r(10) = .714) (Fig. 2I). Lastly, the *Id2* expression profiles of layer 6 were also highly correlated between mice and voles (r(10) = .97), with bootstrapped 95% confidence intervals overlapping for nearly the entire medial-lateral length of the section, including a decrease in expression between visual and auditory cortical areas (Fig. 2L, black arrow).

To adjust for the possibility of inflated or spurious correlations, we normalized expression levels between species by differencing the data along the medial-lateral axis, using percentage change to the previous value instead of absolute magnitude. This cautious approach corrected for differences that may still exist between species even after mean-centering (fig. S2). Correlations of percentage change showed weaker, but still strong correlations between species across cortical layers (fig. S2, right). Posterior layer 2/3 *Id2* expression profiles remained positively correlated between voles and mice (r_% change_(10) = .57). Similarly, layer 5 and 6 *Id2* expression remained significantly correlated (Layer 5: r_% change_(10) = .697), Layer 6: r_% change_(10) = .414). Thus, after controlling for changes in absolute magnitude, expression profile data from caudal cortex confirms that *Id2* expression is tightly conserved on a layer-by-layer basis between mice and voles at early stages of cortical development.

Taken together, between-species correlations show that the spatial pattern and level of *Id2* is moderately conserved across the neocortex in mice and voles, but the extent of conservation varies by layer and area. Notably, while whole-map correlations between species ranged from high to moderate between layers, expression profiles of caudal cortex encompassing visual and auditory cortex specifically, were tightly correlated, suggesting that *Id2* expression in caudal areas is highly conserved. However, the above data also shows that key differences exist between species in layer 6 and rostral neocortex, where the weakest correlations were present. These differences may be due to the fact that laboratory mice are highly inbred and not reflective of their natural counterparts, or they may represent species specializations and changes in developmental trajectories that have emerged in the 35 million years of independent evolution that mice and voles have undergone (*21*).

## Divergent areas of cortical gene expression across species

After finding that *Id2* was conserved between mice and voles across cortical layers, we sought to quantify the extent to which area-specific patterns of expression for *RZRβ*, a long-studied primary sensory area marker, diverge. We hybridized neonatal vole and mouse brain tissue for *RZRβ*, whose high layer 4 expression is heavily influenced by input from thalamic sensory nuclei (*9, 22*). Because *RZRβ* is highly expressed in this layer, and sparse in other layers at early stages of development, we restricted our analysis to layer 4 (Fig. 3, A-D) (*23*). We analyzed serial 100µM sections of tissue, and again reconstructed the data in 3D, digitally unwrapped and flattened the expression patterns, and projected individual patterns into a common reference frame to enable point-by-point comparisons. We then calculated correlations between the transformed RZRβ expression patterns within and across species (Fig. 3A). Vole *RZRβ* expression patterns were highly correlated across individuals (r_avg_(21) = .742 ± .110).

**Fig. 3.**
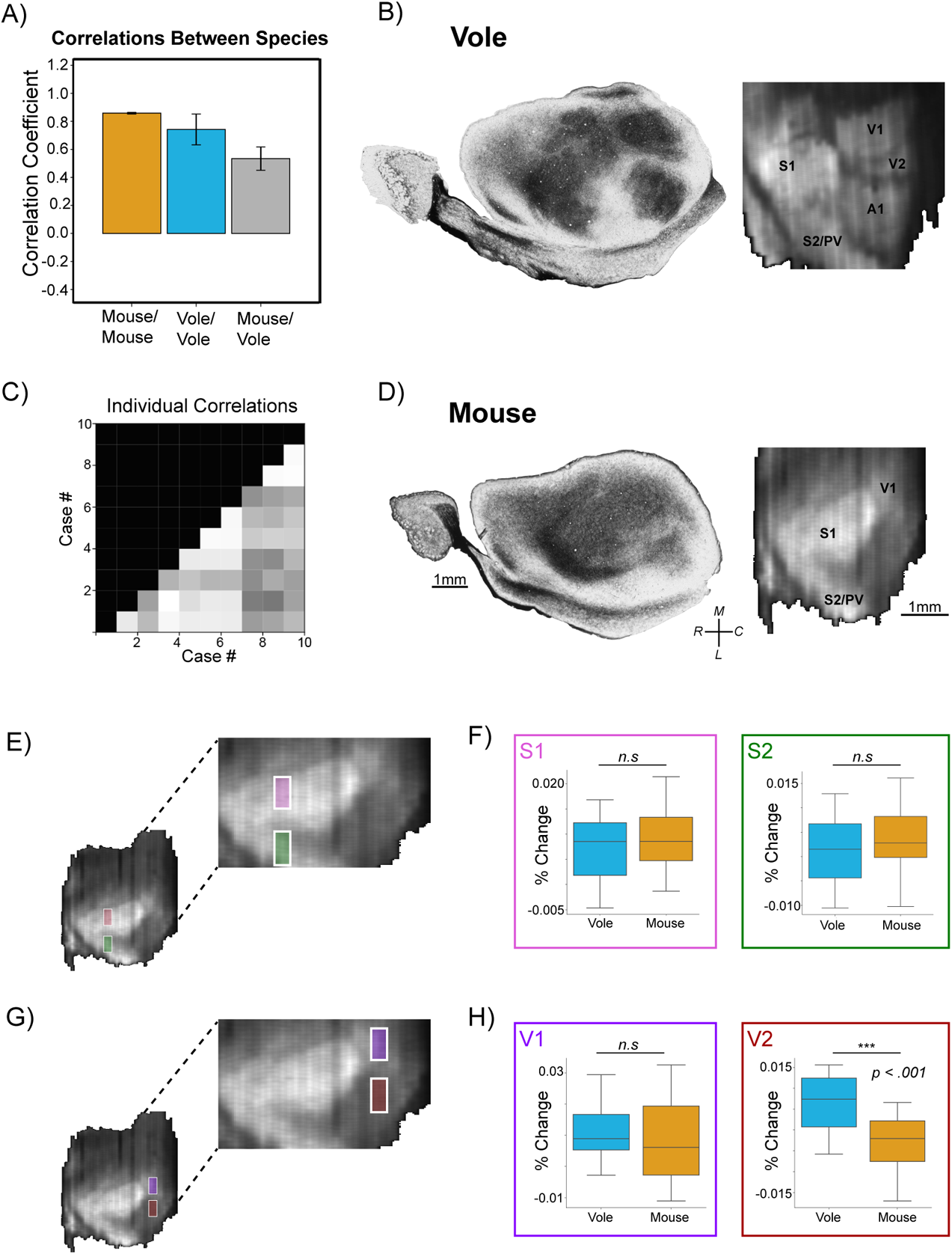
*RZRβ* expression shows divergent patterns for visual and auditory areas between species. (A/C) Bar graph (A) of average correlation coefficients of whole-cortex layer 4 *RZRβ* maps within and across species (vole: n=8, mouse: n=3) shown as mean with standard deviation. Note the low variation between individual mouse maps. Correlation matrix (C) showing correlations between individual cases (White = 0, Dark Grey = 1). (B/D; Left) Flattened cortical sections stained for myelin in an adult vole (B; top) and mouse (D; bottom) showing cortical field boundaries. (B/D; Right) Example cases of digitally flattened *RZRβ* maps reconstructed by Stalefish from coronally-sectioned tissue in a vole (top) and mouse (bottom). The medial wall was cropped in these reconstructions to align data with myelin stains. The pattern of *RZRβ* in layer 4 appears to be coincident with the boundaries of cortical fields in both species. Expression level is represented as pixel-value at a given coordinate (black = lowest, white = highest). (E) Example of a mouse *RZRβ* reconstructed expression map showing the location and relative size of digital micro-punches from S1 (pink) and S2/PV (green). 200µM by 500µM digital micro-punches are drawn to scale on flattened expression maps (post-transform). (F) Box plots of digital micro-punches taken from locations in S1 (pink) and S2/PV (green). In this case, the Y-axis denotes average percent-change (not raw signal level); x-axis denotes species. Box size represents quartiles 1 and 3 with median. Whiskers are set as 2 times the interquartile range. (G) Example of a mouse *RZRβ* expression map showing the location of digital micro-punches from V1 (purple) and V2 (red). (H) Box plots of digital micro-punches taken from V1 (purple) and V2 (red) with the same parameters as F. Abbreviations: Primary Visual Cortex (V1). Secondary Visual Cortex (V2). Primary Somatosensory Cortex (S1). Secondary Somatosensory Cortex (S2). Parietal Ventral Area (PV). Not significant, p > 0.05 (n.s). p < .001 (***).

Remarkably, not only were mouse *RZRβ* maps highly correlated with one another (and significantly more so than with voles: p = .009), but the amount of individual variation was near zero (r_avg_(3) = .858 ± .006). As with *Id2*, cortex-wide *RZRβ* expression patterns were well correlated between species (r_avg_(21) = .534 ± .083). These results were expected given the evolutionary and life history of these rodent species, and tightly regulated thalamocortical development, which influences *RZRβ* expression. However, the moderate (not strong) correlation in expression between species indicates also that differences have emerged in each lineage.

Because *RZRβ* was found to be expressed in layer 4 in both species and at similar levels, we hypothesized that species differ in terms of the identity of the putative cortical area in which *RZRβ* is expressed, rather than in terms of cortical layer. To begin to address where these differences may have emerged, we compared reconstructed neonatal *RZRβ* cortical expression maps to flattened cortical sections of adult animals stained for myelin, which show clear delineations of cortical field boundaries. We found that *RZRβ* expression in the early postnatal brains revealed the putative cortical areas that were later delineated by myelin stains in adult neocortex (Fig. 3, B and D) (*24, 25*). Specifically, reconstructed *RZRβ* expression patterns clearly demarcated the cortical field boundaries of putative S1, A1 and V1 in the prairie vole (Fig. 3B). While expression patterns for neonatal mice revealed a clearly defined primary somatosensory cortex (Fig. 3D), they did not reveal the putative A1 and V1 to the same extent. In both species, the sub-regions representing the map of the entire body could be identified from the expression patterns in putative S1 (*26, 27*). To our surprise, the second somatosensory/parietal ventral area (S2/PV) was also delineated in both species. Both mice and voles also displayed *RZRβ* expression in V1, but this expression was more clearly defined in prairie voles. Furthermore, across all vole cases (n = 7), *RZRβ* expression patterns displayed a distinct boundary between the primary and the second visual areas (V1 and V2). To our knowledge, evidence for distinct higher order (HO) cortical fields has not previously been demonstrated at such an early stage in development, and this result suggests that thalamocortical interactions or early arealization gradients may play a stronger role in patterning HO areas than has been previously thought (*28*). However, we also note that mice lacked a clear delineation between these visual areas and displayed an absence of *RZRβ* expression in putative A1.

We next quantified the magnitude of these species differences by taking digital micro-punches of regions of interest, whose location was determined post-hoc, i.e., after examining cortex-wide difference maps. These maps were created by subtracting the average mouse expression map from the average vole expression map (data not shown). We selected micro-punch regions of interest corresponding to peaks in expression of putative S1, S2/PV, V1, V2, and A1 (Fig. 3, E-H). We then compared the average post-differencing signal value around each expression peak, finding no significant difference in the change in peak expression between species in S1 (anterior barrel cortex) (Fig. 3F; pink box) or S2/PV (Fig. 3F; green box) (S1: R^2^_adj_ = .023, F(2) = 1.458, p = 0.255, S2/PV: R^2^_adj_ = −.019, F(2) = .636, p = 0.267). While less apparent when observing average or difference maps, we also found no significant difference in the average percentage-change in expression level between species in V1 (Fig. 3H; purple box) (R^2^_adj_ = .091, F(2) = 2.961, p = 0.442), which means that for this portion of V1, voles and mice showed similar changes in levels of expression. On the other hand, neonatal voles had significantly higher changes in *RZRβ* expression in V2 than neonatal mice (Fig. 3H; red box) (R^2^_adj_ = .315, F(2) = 9.977, p < 0.001). Neonatal voles also showed increased *RZRβ* expression at the spatial location for A1 (R^2^_adj_ = .382, F(2) = 9.037, p = 0.013) (data not shown). Thus, comparison of digital micro-punches from putative cortical fields across species suggests that layer 4 *RZRβ* expression is tightly conserved for some areas (S1, S2/PV, V1), and not others (V2, A1). Together with the correlational data, we interpret these results as evidence that spatial expression patterns and signal levels of *RZRβ* expression are conserved across the somatosensory cortices of mice and prairie voles, but that the patterns and levels of *RZRβ* expression in visual and auditory areas vary between species. Similar to *Id2*, these differences may be products of independent evolution and associated with species specific behaviors mediated by these different sensory systems, or to changes in mice due to inbreeding.

## Conclusions

Here we present a new open-source software tool, *Stalefish*, which allows for the rapid acquisition and analysis of ISH data, and quantification of cross-species comparisons by projecting reconstructed data into a common reference frame. Our results indicate that cortex-wide *Id2* expression patterns in the neocortex of early postnatal brains are conserved between species in a layer-dependent manner, ranging from moderately (Layer 6) to highly (Layers 2/3 and 5) conserved. However, in caudal neocortex, all layers yielded strong correlations, suggesting that species-specific expression of *Id2* is restricted to rostral areas. This analysis highlights the usefulness of *Stalefish* for analyzing whole brain regions, for observing overall patterns, and for allowing the scientist to then focus on ROI’s in search of the factors that drive strong and weak correlations, i.e., the factors underlying species similarities and differences. Specifically, the *Stalefish* tools enabled us to observe that *RZRβ* is expressed in higher-order cortical areas such as S2/PV and V2. This discovery would not have been possible with traditional analysis.

It would next be of great interest to catalogue the emergence of these HO areas along with how other patterns of expression in the neocortex change across the first two postnatal weeks, to study species or experimental differences in developmental trajectories in quantitative terms and at the level of entire neural structures. The *Stalefish* algorithms and software tools are readily applicable for the analyses of multiple species over multiple postnatal days, to elucidate where and when developmental processes are conserved or have diverged in evolution. Further, they allow for the study of how variation emerges across development and across species. Lastly, while we used *Stalefish* to show how spatial patterns of *Id2* and *RZRβ* are conserved in the cortex, these tools can easily be applied to study laminar expression profiles (fig. S3), or other brain structures whose spatial patterns of gene expression have eluded quantitative study for decades, such as the hippocampus and dorsal thalamus (fig. S4 and S5). While we present the first data using this new methodology in the present study, we believe that neuroscientists will be able to utilize *Stalefish* to quantify spatial patterns of gene expression (or histological markers) in a variety of brains of different shapes, sizes and levels of complexity, and be able to build on this approach to address longstanding evolutionary and developmental questions, generating novel comparisons and deriving unique insights that may otherwise have remained elusive.

## Acknowledgments

Thank you to Dr. Andrew Fox and Dr. Andrew Halley for initial comments on conceptualization of the software.

## Author contributions

Conceptualization: ME, SJ Methodology: ME, SJ Investigation: ME, SJ, RB, SW Visualization: SFB, MJM, JLS, EH Funding acquisition: LK, SW Supervision: LK, SW

Writing – original draft: ME, LK, SW, SJ

Writing – review & editing: ME, LK, SW, SJ, KH, RB

## Data and materials availability

All data are available in the main text or the supplementary materials, and at https://github.com/ABRG-Models/Stalefish.

## Supplementary Materials

Materials and Methods Supplementary Text, Figs. S1 to S14 Tables S1 to S3 Movies S1

## Supplementary Materials

### Materials and Methods

#### Subjects

Fifteen prairie voles (*Microtus ochrogaster*) and six C57/BL6 Mice (*Mus musculus*) were used for In-Situ Hybridization (ISH) experiments. Voles were obtained through the breeding colony at the University of California Davis, and mice were obtained through the breeding colony at the University of California Riverside. All experimental procedures were approved by UC Davis IACUC and UC Riverside IACUC and conform to NIH guidelines.

#### Brain collection

Tissue was collected on postnatal day 1 for voles and postnatal day 0 for mice. Animals were euthanized by an overdose of Sodium pentobarbital (> 100 mg/kg, 390 mg/ml) and perfused with 0.1 M phosphate buffered saline (PBS) followed by 4% paraformaldehyde (PFA) in 0.1 M phosphate buffer. Brains were extracted under microscope guidance and stored in 4% PFA before being shipped to UC Riverside for processing. Brains were then dehydrated in ascending concentrations of methanol and stored in 100% methanol at −20 degrees. Brains were fixed in gelatin-albumin solution and sliced on a vibratome at 100um. Alignment landmarks for use in post-processing were created by positioning a straight 21-gauge needle through the mold in which brains were fixed in the gelatin-albumin solution (dissolved in 1X phosphate buffered saline and fixed with 25% glutaraldehyde). Needles were removed once the gelatin-albumin fixing medium had solidified, leaving circular holes in each slice that aid the subsequent alignment process (see *Supplemental Methods*, Figure S1).

#### In-Situ Hybridization

Previously established protocols for non-radioactive free-floating RNA in-situ hybridization (ISH) were used to assess patterns of gene expression in mice and voles ^19, 29^. Probes for *RZRβ* and *Id2* were applied to alternating sections of 100μm coronal slices. After hybridization, sections were permeabilized in 50% glycerol, mounted onto glass slides, and cover-slipped. All hybridized sections were digitally imaged using a Zeiss SteREO Discovery V.12 dissecting microscope and captured using a digital high-resolution Zeiss Axio camera (HRm) using Axiovision software (version 4.7).

#### Overview of 3D reconstruction, digital unwrapping, and point-by-point alignment

In general terms, the 3D reconstruction process consisted of i) fitting curves to the elements of a common anatomical surface that are visible across multiple 2D slice images; ii) sampling image luminance values in contiguous rectangular bins oriented tangential to each curve at evenly spaced points along its length; and iii) aligning the data obtained with respect to each curve to form a 3D surface that corresponds to the shape of the original anatomical surface from which the curves were derived. The ‘pre-sliced’ alignment within the stack of slice images was then approximated using an algorithm that aligns each curve with respect to the curve obtained from the adjacent slice, utilizing, if necessary, any available alignment marks e.g., circular holes left by a needle. Each 3D surface was then digitally unwrapped with respect to a user defined ‘brain axis’ and an angle about this axis which formed a center line through the surface (see Fig. 1 and Supplemental Methods), allowing the curves to be digitally straightened while clamped to the center line. Finally, each resulting 2D expression map was linearly transformed so that its coordinates matched those of a comparable reference map obtained from another animal. This was achieved by marking three external anatomically-defined locations for each structure near to the curve surface elements on selected brain slices and then, following reconstruction and digital flattening, finding the 2D linear transform to project this triplet of coordinates onto the corresponding triplet on the reference map. The linear transform was then applied to all of the sampling bin locations in the dataset for each individual, resulting in a map for each that was composed of irregularly sized, quadrilateral data pixels. These data were then resampled, using elliptical Gaussian kernels, onto a Cartesian grid of square pixels, to form a 2D matrix of binned luminance values that could be compared to similarly resampled data from the reference map on a point-by-point basis.

#### Curve fitting and image sampling

To allow the researcher to define arbitrarily shaped, smooth curves that follow anatomical feature lines such as those shown in Fig. 1, B-D, we utilized the Bezier curve, a form of polynomial curve commonly used in drawing software, which is typically defined by a start and end location and a series of user-editable ‘control points’ that define its curvature. While these control points (which typically lie away from the curve) give great flexibility for drawing applications, we designed the *Stalefish* curve drawing tool to exploit the fact that it is also possible to analytically determine a Bezier curve that best fits a given sequence of points. This allowed us to mark points along the boundary of an anatomical structure in each slice without considering where the Bezier control points should lie. The number of ‘user points’ in the sequence determined the order of the polynomial which formed the Bezier curve, with three points specifying a quartic curve, four specifying a cubic, and n+1 points in general specifying an n^th^ order curve. Fits of the highest quality were obtained for our data by using multiple, low-order curves, joined end-to-end using a simple routine to modify the control points closest to the join to match the gradient at the end of one to the gradient at the start of the other. Once the (multi-segmented) curves were defined by the user, N sample boxes were automatically defined by drawing N+1 equally-spaced vectors along the curve and normal to it. To find N+1 equally spaced locations on the curve, the following three-step procedure was carried out numerically: First, the distance between the first and last points on the curve was computed and divided by N to get a candidate spacing, *s*. Second, up to N times, a Euclidean distance was advanced *s* along the curve, recording the coordinate at each step. Noting that Bezier curves are parameterized with *t* in the range [0,1] (mapping coordinates on the curve from start to end), the increment of *t* which advanced a coordinate a distance *s* along the curve was computed via a simple binary search. The algorithm accounted for the steps that crossed the join of two Bezier curves. Third, the number of coordinates that could fit onto the full curve for spacing *s* was reviewed, and if that was different from N+1, *s* was adjusted (by doubling/halving it) and the second step repeated, until the number of coordinates on the curve was N+1. The start and end of adjacent normal vectors provided four corners of a box from which pixel intensities were sampled. These methods were used to measure the variation in average signal intensity along one or more anatomically aligned curves identified in each slice image.

#### Slice alignment

To assist the process of aligning curves from consecutive slice images, a needle was used to create visible markers in all slices corresponding to a given brain (see fig. S6). Then, on the image of each slice three user-defined points were digitally marked on the perimeter of the needle hole, from which the parameters of a circumcircle were calculated to estimate the center of the hole. A two-dimensional coordinate offset was then applied to the data sampled from each slice to place the alignment landmarks on an ‘alignment axis’ in 3D space that we defined, by way of convention, to be parallel with the x-axis. Then, starting with the second slice image, each image was automatically rotated about the alignment axis so that the points on the curve were as close as possible to the points on the curve in the previous slice. The optimum rotations were determined by minimizing the sum of squared distances between N equally spaced locations on the curve on slice *i* and the corresponding N locations on the curve of slice *i*-1.

#### Digital unwrapping

Digital unwrapping is the process of straightening out a curved, three-dimensional surface into a two-dimensional map. This process began with a set of aligned curves (see *Supplemental Methods*, fig. S12 A). We placed axis marks that defined a brain axis (white bars in fig. S12). An unwrapping axis of ‘zero marks’ was defined on the surface, by rotating a user-defined angle about the x-axis (centered on the brain axis), then locating the most distal point on each curve at this angle (blue/rainbow-colored spheres in fig. S12 A). Each expression ‘ribbon’ was then straightened out, keeping it fixed at its zero mark (fig. S12 B). The final step was to resample the image in Figure S8 E to produce an image consisting of square pixels, as shown in fig. S12 F.

#### Digital reconstruction of *Id2* and *RZRβ* expression patterns

For each gene/layer, curves of the cortex were semi-automatically traced using the *Stalefish* software tools (See *Supplementary Text* and Video S1). For optimal resolution, we chose 150 bins for data collection, which spanned medial to lateral. For each slice, data collection began at the most medial aspect of the medial wall (near the subiculum) and continued laterally to the rhinal fissure. While we used standard ISH, fluorescent or multi-colored ISH are also easily analyzed by *Stalefish*. To align slices we used either the *Stalefish* landmark alignment mode (where possible) or the circle-mark mode in combination with placing axis-mark data-points at the beginning and end of each brain (See *Supplementary Methods*). To place brains of different individuals or species into a common reference frame for point-by-point comparison, we placed three landmarks at the same morphological location in each case, e.g., with one at the apex of medial wall 200µM in the slice adjacent to that in which hippocampal formation was first visible. 200µM expression profiles were taken by simply retrieving the pixel value and location data stored in the h5 file of each map and selecting columns of the data-frame which corresponded to the region of interest (post-transform). Similarly, digital micro-punches (used in Figure 3) were taken in a similar manner but used both rows and columns (500µM by 200µM). Thus, the output data from *Stalefish* is easily selectable, choosing any span of data or ROI, simply by indexing the output data-frame.

#### Statistical analysis

Correlations were calculated using Pearson’s correlation coefficient; r_avg_ is reported with the number of correlations used to generate the average correlation coefficient ± standard deviation, and r is reported as r(*n*), where *n* is equal to the number of subjects. To assess between-species correlations, all maps of individual cases were transformed using the above method to a single case. Data was differenced when necessary, using the percentage change differencing method within Numpy and SciPy Python packages. In Figure 3, boxplots are presented with quartiles 1 and 3, the median, and whiskers corresponding to 2 times the interquartile range. Boxplots were created using average percent-change and not raw signal, to show statistical differences in the *change* in signal. Statistical tests for correlations and boxplots were conducted using linear models, where distance along the cortex was used as an interaction term when appropriate. The general form of these equations used species and location as predictors for signal level, correlation, or percent change. The results of these tests are presented with the adjusted R-squared value (R^2^_adj_) and F-statistic F(*m)*, where *m* is equal to the number of degrees of freedom of the model.

**Fig. S1.**
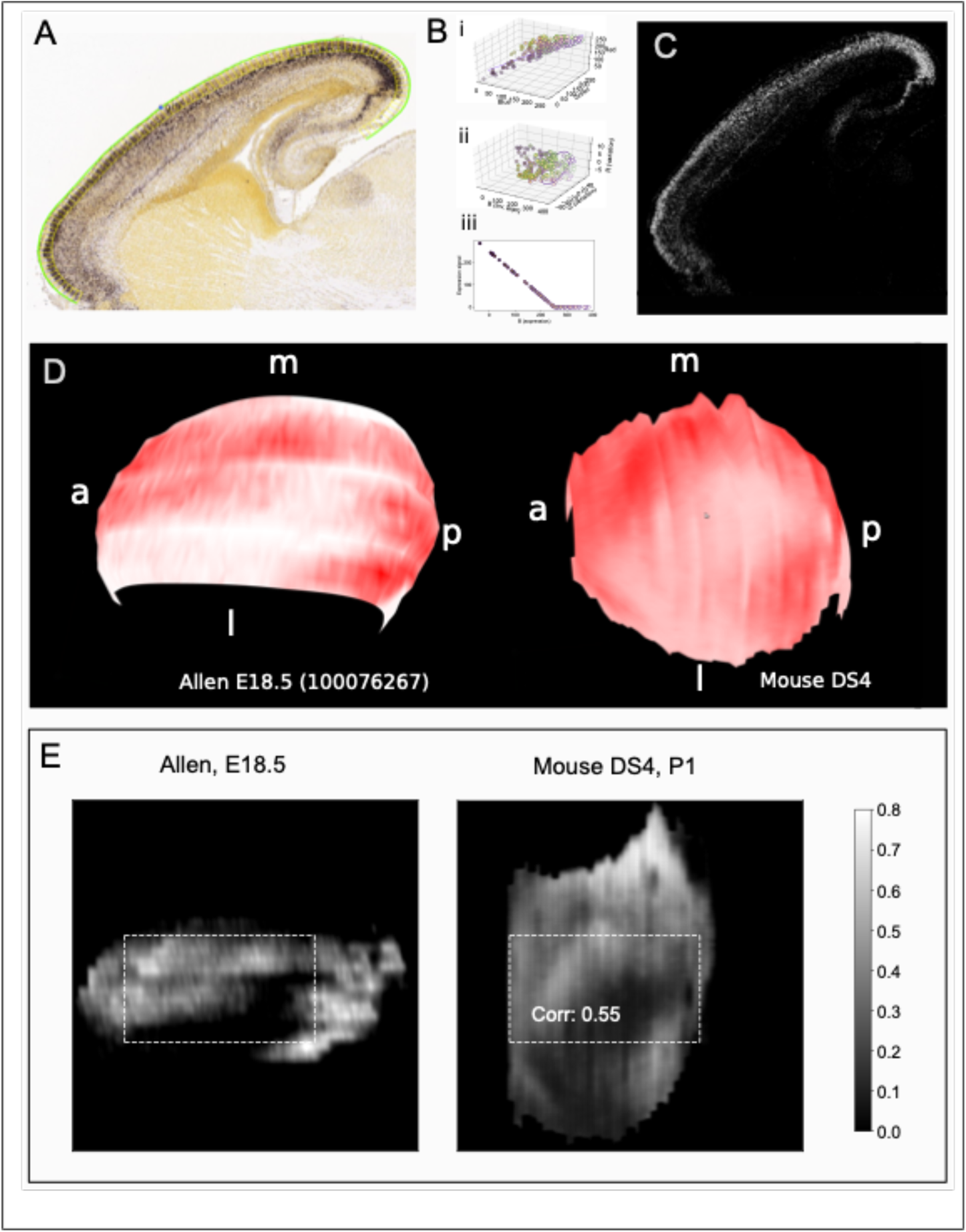
*Id2* gene expression reconstructed from Allen Developing Mouse brain slice data. (A) Allen ISH images are colored and sagitally sliced, in contrast to our data which are monochrome stains of coronally sliced brains. In the Allen data, dark purple indicates *Id2* gene expression. (B) Extracting the signal from the image. The Allen images are provided with an associated image which shows ‘expressing pixels’ in a colormap but without information allowing the expression levels to be converted to a number. We set about determining a technique to convert from color values in the original images to a signal value. (i) Pixels plotted in red-green-blue 3D space. Allen-specified non-expressing pixels have a green outline, expressing pixels have a purple outline. We made a linear fit to the expressing pixels. (ii) To achieve this, we rotated the data in color space until the linear fit through the expressing pixels lies on the ‘blue’ axis. We allowed some variation in color about the fit by encircling the axis with an ellipse. Pixels falling within this elliptical tube were deemed to be ‘expressing’. (iii) The expression level is inversely proportional to the brightness of the pixel and we set a cut-off brightness above which the expression is set to 0. (C) The resulting signal corresponding to panel A. (D) Three dimensional Layer II-III expression surfaces for the Allen data (left) and our data. Key: a: anterior, p: posterior, m: medial, l: lateral. (E) Digitally unwrapped surfaces generated from the 3D data in D. Both datasets have the same anatomically determined landmarks and the Allen data has been linearly transformed to match our data, so length scales are unified. The Allen dataset is partial (some lateral slices are missing) and so the Allen map appears ‘wide and narrow’. The dotted rectangle marks the region for which both maps have data. Visual inspection of the content of the rectangles suggests that there is good correlation between the images, with a dark region of low expression from the bottom left to the middle right apparent in both maps. A Pearson correlation of the pixel values within the rectangle of 0.42 lends support for this interpretation. References: *Lein, E.S. et al. (2007) Genome-wide atlas of gene expression in the adult mouse brain, Nature 445: 168-176. doi:10.1038/nature05453*. Image used for E18.5 Id2 experiment: https://developingmouse.brain-map.org/experiment/show/100076267. Slice shown in panels A and C: http://api.brain-map.org/api/v2/image_download/101267565.

**Fig. S2.**
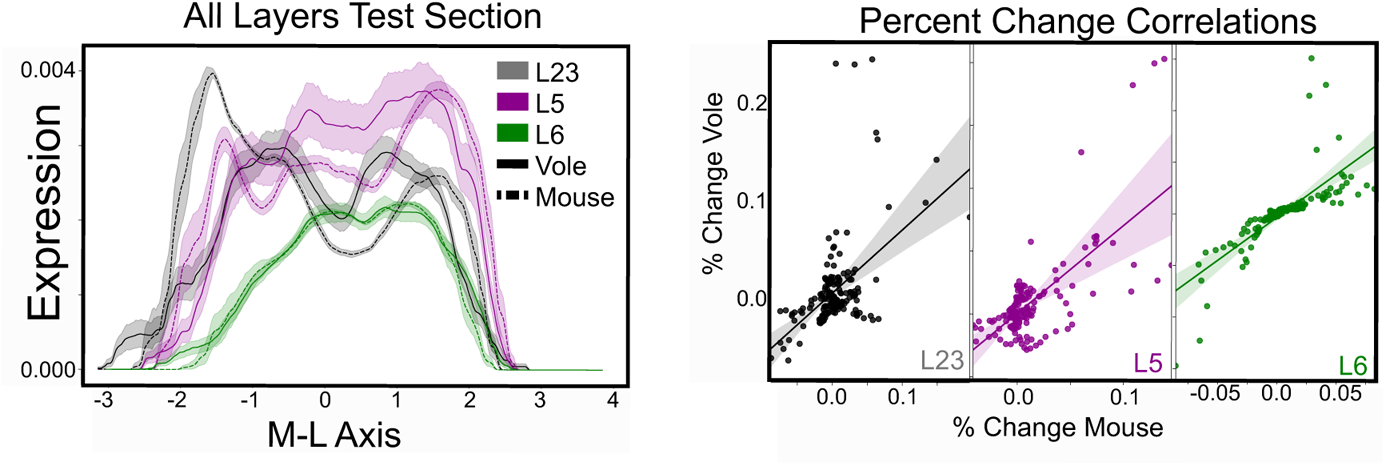
Percent Change Correlations. Line graphs (left) show *Id2* expression profiles of layers through a single 200uM profile taken from post-transform posterior cortex: Layer II/II (L23) gray, Layer V (L5) purple, Layer VI (L6) green. Vole (solid line). Mouse (dotted line). Line graphs are presented as mean with bootstrapped 95% confidence intervals. Scatter plots (right) with line of best fit and 95% confidence intervals show correlations between expression percent change in mice (x-axis) compared to voles (y-axis) at each location along the expression profile of each layer.

**Fig. S3.**
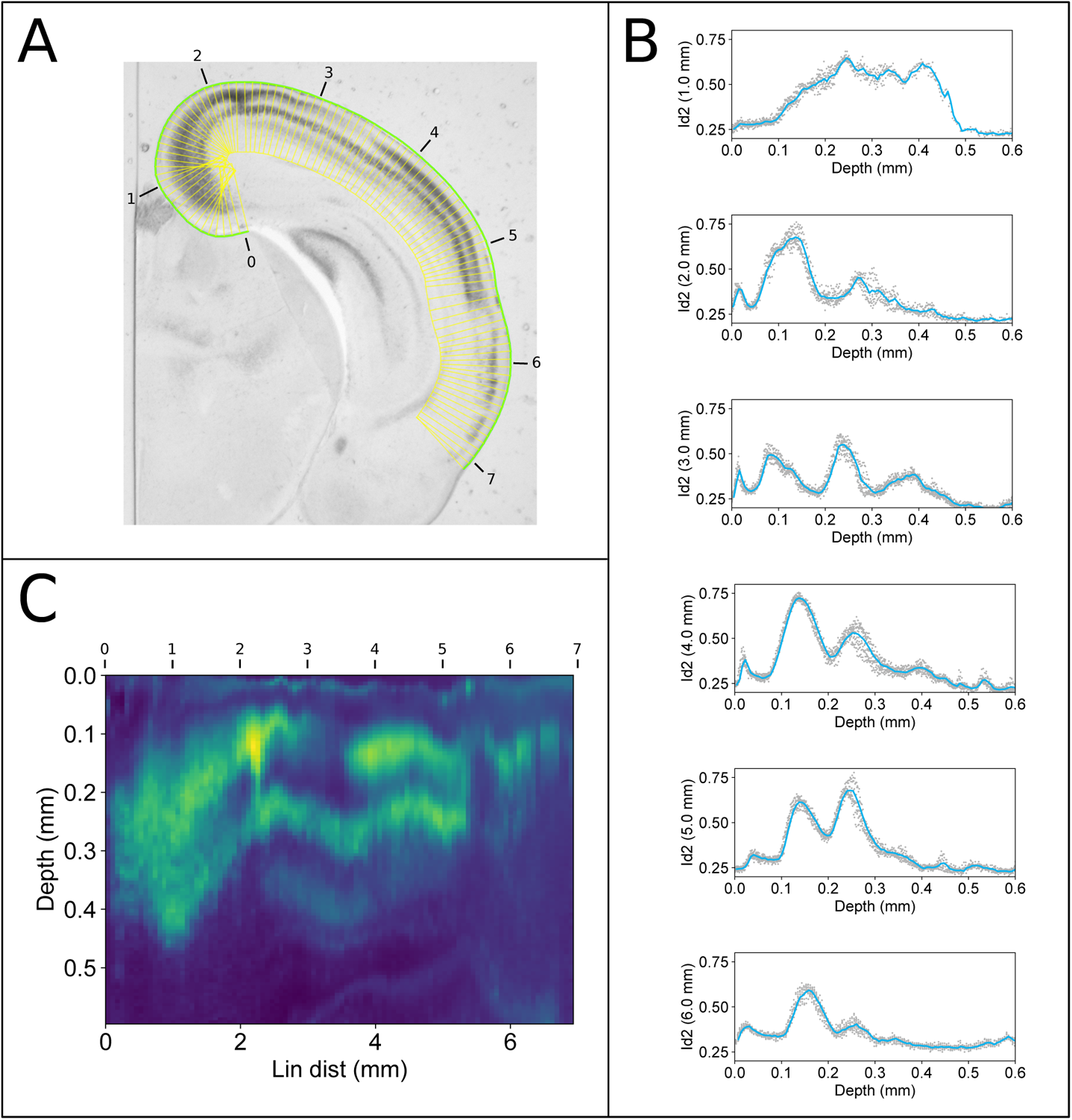
Depth profiles for laminar analysis. (A) User-supplied curve is shown in green, and sampling boxes in yellow. The linear distance along the green curve is marked in black annotations (units are mm). The boxes have depth 0.6 mm. (B) Gene expression as a function of box depth for sample boxes at 1 through 6 mm of linear distance along the green line in A. Grey dots are the individual pixel values of the pixels within the sampling box, the blue line is a histogrammed mean expression (100 bins). (C) A heat map of Id2 gene expression as a function of linear distance along the green curve in A. The depth values are extracted from the blue histogram values show in B for 6 selected linear distances.

**Fig. S4.**
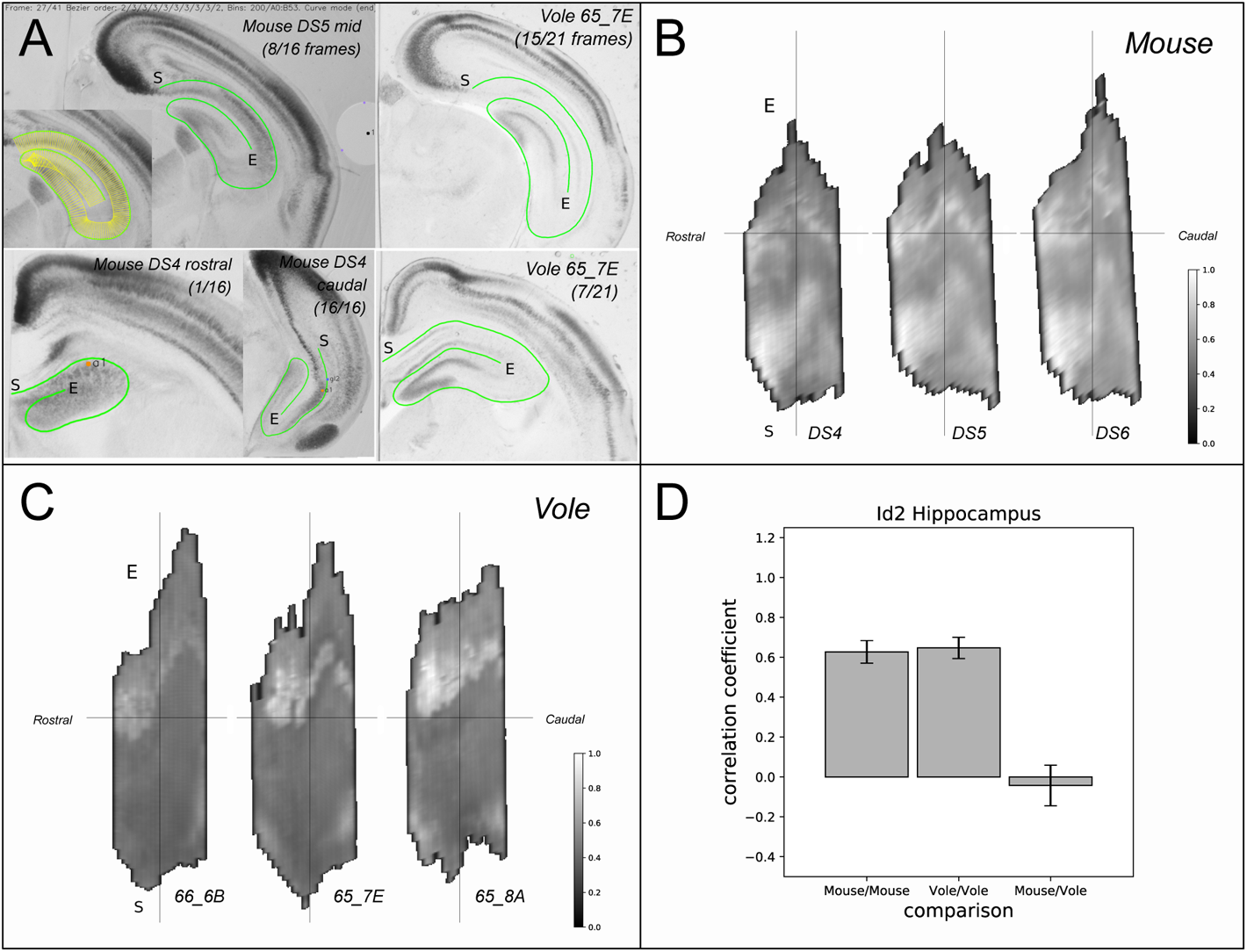
A demonstration of the use of *Stalefish* in an alternative brain structure. Our initial motivation for developing the *Stalefish* technique was to learn about interspecies differences in gene expression in the mammalian neocortex. The ability to examine depth-based expression is ideal for studying layer-specific expression in the neocortex. However, the technique is applicable to any structure in the body and so we applied it to *Id2* gene expression in the spiral structure of the Hippocampus. (A) Spiral sampling curves allow the digital unwrapping of the hippocampus to examine and compare Id2 expression in mouse and vole. Curves were marked out in a clockwise direction; the start is marked with S and the end of each curve is marked with E. Axismarks were carefully chosen (visible the Mouse DS4 examples) to provide ‘zero-angle’ marks along the dorso-lateral Hippocampus. (B) Unwrapped hippocampi for three mouse brains. Each brain was marked with three landmarks, although these were based on gene expression rather than on specific anatomical features. The landmarks have allowed the mouse samples to be transformed onto the coordinate frame of one of the vole brains (66_6B). Assuming that illumination and stain response are similar for each dataset, the signals have been normalized as a group (the colorbar applies to all three maps). The Id2 expression in the dentate gyrus is visible in the top half of the maps; there is also widespread expression in the lower half of each map. (C) Similar unwrapped hippocampi for the vole, transformed onto the coordinate frame of sample 66_6B. The vole hippocampus has strong expression in the dentate gyrus, but minimal expression in the bottom half (CA1) of the maps. (D) Pearson correlation coefficients for mouse to mouse, vole to vole and mouse to vole comparisons show that the maps are well correlated within a species group, but that the expression present in the lower half of the maps for mouse destroy the correlation between species. Error bars are standard deviations for the correlations of all possible map pairs.

**Fig. S5.**
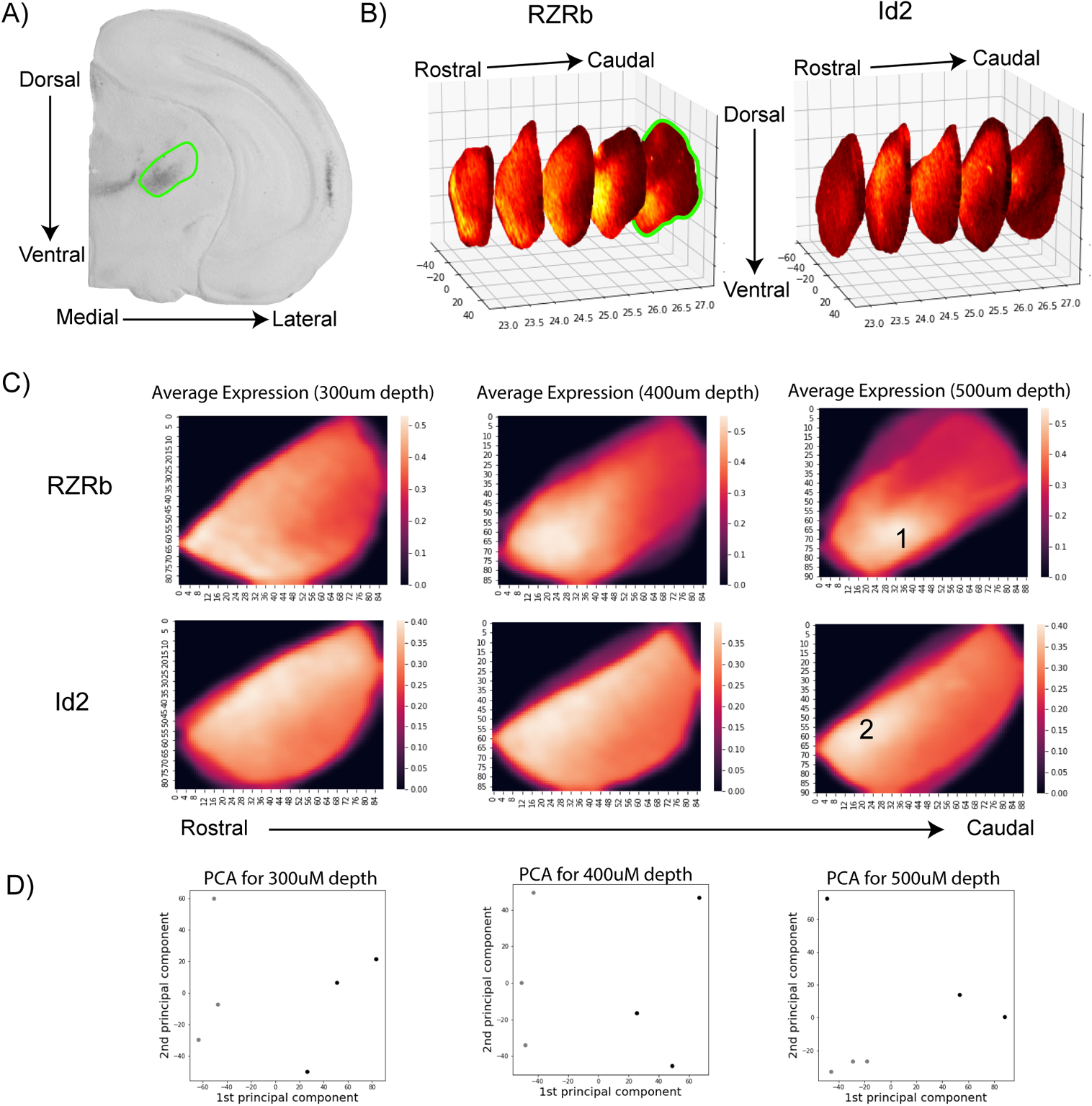
Thalamic gene expression in the ventral posterior nucleus in mouse. (A) Coronal section of vole brain tissue hybridized for *RZR*β mRNA. Green circle denotes which region was analyzed using the freehand tool in *Stalefish*. (B) 3D graph representing various sections (Anterior-Posterior Plane). Green highlighted section is data retrieved from (A). Axes denote the 3 spatial dimensions of the brain. (C) Average expression maps (n=3 per slice), of serial coronal sections (akin to those shown in A, but averaged over multiple cases), for *RZR*β and *Id2*. Note how *RZR*β expression is limited to the lateral portion of VP (black number 1), while Id2 is restricted to the dorsal aspect of VP (black number 2). (D) A simple principal components analysis showing how dimensionality reduction can be used to show the relationship between the spatial expression of *Id2* (black) and *RZR*β (grey).

## Supplementary Text

### Introduction

This document provides an extended description of the Stalefish analysis process described in the main paper and available at https://github.com/ABRG-Models/Stalefish. Because the technique requires no special equipment, it is accessible to any lab already equipped to image histologically processed brain slices. To help other researchers create similar analyses using their own data, in the first section of this document, we present a tutorial style description of the process used to create coordinate-centered expression maps from a set of brain slices along with details of the algorithms used in the program.

The central idea of the process is to fit smooth, anatomically relevant curves to the structures visible in the 2D slices, sampling the image luminance in the region below the curve at equally spaced locations along its length. These curved sets of ‘luminances below the curve’ for each of the slices are then joined together so that a 3D surface is created. An algorithm makes an approximation to the ‘pre-sliced’ alignment of the individual slice images, by aligning each curve with respect to its neighbor (the best alignment is achieved if, during the experimental procedure a visible alignment mark is made by inserting a needle through the brain mount material). The 3D surface is then ‘digitally unwrapped’. The researcher defines a ‘brain axis’ and an angle about this axis which forms a ‘center line’ through the surface. The curves are ‘digitally straightened’, each one being clamped to the center line. The resulting 2D map can be linearly transformed so that its coordinates match those of another brain. This is achieved by marking three anatomically identifiable locations on each brain, ideally close to the curve surfaces. The linear transform required to transform one triplet of coordinates into the template coordinates is computed and then applied also to the luminance ‘data pixel’ coordinates. Finally, the map of irregularly sized, quadrilateral data pixels is resampled onto a Cartesian grid of square pixels making a regular, quantitative image which is easy to submit to standard point-by-point analysis methods.

Although we present this technique as it is applied in the main paper to in-situ hybridization (ISH) stains for the genes *Id2* and *RZRB*, it could be applied to any stain in which there is a reliable, monotonic relationship between image luminance and the value of a variable of interest. For example, this technique could be applied to cytochrome oxidase or nissl stains. Although the data presented in this work is based on monochrome ISH images, the software can also be used to interpret colored stains. As a demonstration, we include 3D reconstructions of data from the Allen Developing Mouse Brain Atlas.

In addition to a detailed description of the process in the first two sections, we provide here a step-by-step protocol for capturing expression surfaces from slice sets, and we present a number of additional analyses and visualizations to demonstrate what can be achieved once a set of expression maps have been transformed onto a common coordinate system.

### Sample preparation and image capture

Samples are prepared in a conventional manner, with brains (Fig S6 A) fixed in gelatin-albumin (Fig S6 B) and sliced on a vibratome. The one extension to the usual method is to optionally introduce ‘alignment landmarks’ into the samples. This was achieved by positioning a straight, 21-gauge needle through the mold in which brains were fixed in the gelatin-albumin solution (dissolved in 1X phosphate buffered saline and fixed with 25% glutaraldehyde). When the gelatin-albumin fixing medium solidified, the needle was removed which resulted in the circular marks (visible in Fig S6 C), which shows the resulting brain slice images. This sequence of images serves to illustrate the wide range of shapes formed by the brain as the slices are viewed in a rostral (left) to caudal (right) progression.

**Fig. S6.**
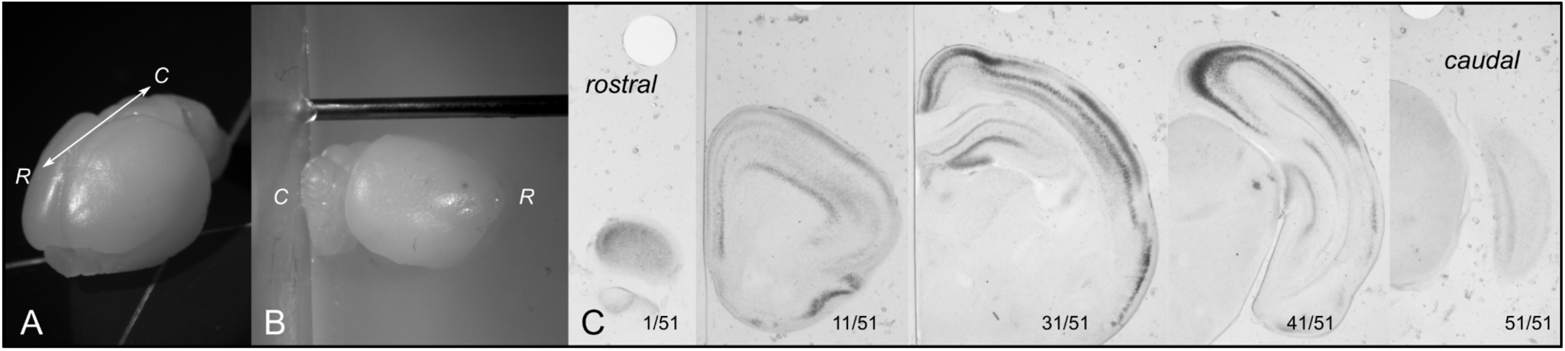
(A) whole vole brain (B) The brain is positioned ready to be set in gelatin-albumin solution, with a needle in place to define the circular alignment landmarks (C) a selection of the 51 coronal sections into which another vole brain was sliced illustrating the diversity of structural shapes of the cortical region.

### Bezier curves

To allow the researcher to define arbitrarily shaped, smooth curves that follow anatomical feature lines such as those shown in Fig. 1, B-D (main paper), we employed Bezier curves; a form of polynomial curve. Commonly used in drawing software, Bezier curves are typically defined by start and end locations and a series of user-editable ‘control points’ that lie away from the curve and determine the curvature. However, it’s also possible to define a Bezier curve that best fits a sequence of points. The control points still exist but are analytically determined from the points, and thus can be assigned automatically once the user has identified several points along the edge of an anatomical feature. The number of ‘user points’ in the sequence determines the order of the polynomial which forms the Bezier curve. Three points gives a quartic curve; four give a cubic and N+1 points give an N^th^ order curve. In principle, an unlimited number of user points could be placed along an anatomical structure and an N^th^ order Bezier curve fitted to them. In practice, the Bezier curve suffers from overfitting for N greater than about 5, and the curve becomes ‘wobbly’, passing exactly through the N+1 user points, but failing to follow the smooth curve of the structure (Fig. S7 A). However, a low order polynomial curve is limited in the complexity of the curve it can fit (Fig S7 B). The most effective curve fitting is achieved for low order polynomials applied to ‘short sections’ of an overall curve as in Fig. S7 C. This fails to fulfil our need to fit smooth curves to structures with complex shapes, such as the hippocampus shown in Fig S7.

**Fig. S7.**
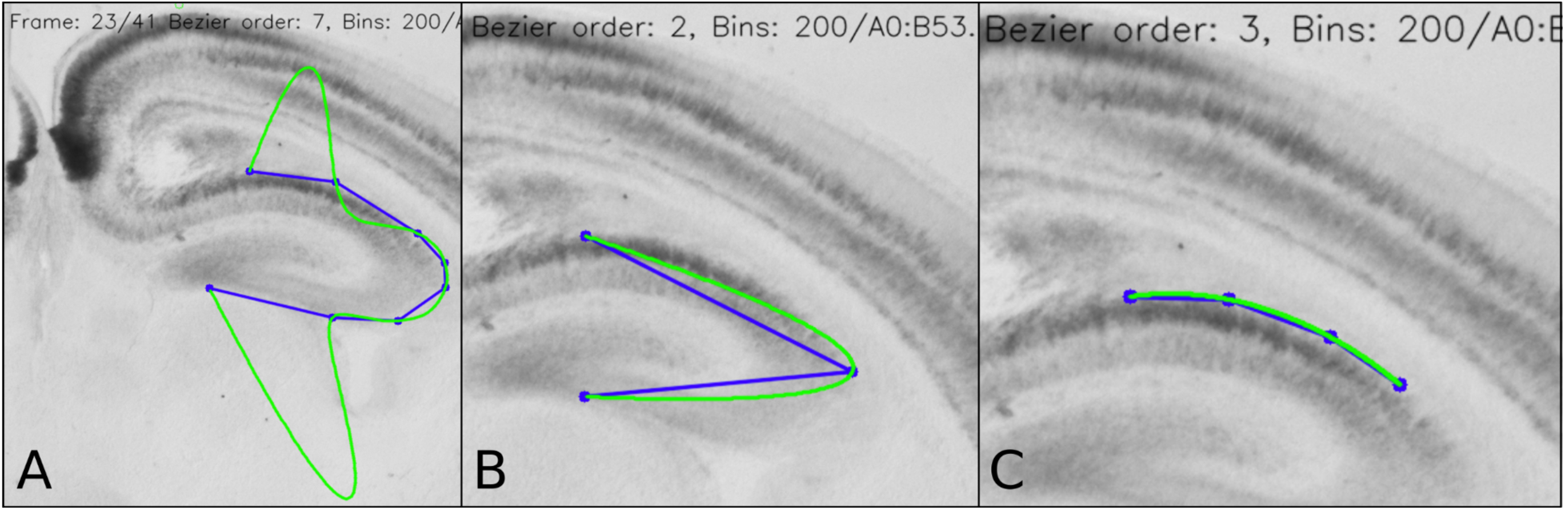
(A) This 7th order Bezier curve demonstrates the problem of overfitting. The curve passes through each of the blue points perfectly, but fails to follow the real shape of the structure that the points are marking out. (B) The converse issue of underfitting, where a 2nd order curve cannot reproduce the curve around the Hippocampus. (C) A 3rd order curve fits a shorter section of the hippocampal curve.

One way to fit curves to complex structures while avoiding overfitting is to allow the user to chain several separate Bezier curves together, with each curve spanning a section of the structure that is short and ‘uneventful’ enough to be fit by a low-order polynomial. However, this presents the immediate problem that two curves joined together at a common point are not guaranteed to have an identical gradient at the join. As we wish to sample from boxes which extend along the normal to the curve, this would lead to non-parallel sampling boxes at the joins. To join the Bezier sub-curves, and provide an overall smooth curve with no discontinuities in its gradient, we used a simple algorithm which modifies the control points closest to the join of the two Bezier curve segments in order that the gradient at the end of one matches the gradient at the start of the next. The algorithm is described visually in Fig. S8. In practice, the researcher adds two or three new points along the curve of the structure she is tracing (we did not fix the order of the individual ‘sub-curves’, allowing the user to experiment), presses a key to ‘commit’ the curve, then adds a few more points for another curve, repeating the process until the entire structure has been traced. It does not always produce an excellent result at the first attempt, but by cancelling and re-drawing points that express the curve of the structure, the researcher can quickly find a good fit. We have found this to be effective and straightforward enough to allow a structure to be traced across a set of 50 slices within about one hour. In future work, it may be possible to further optimize the modification of the control points at the join, to minimize the deviation of the modified curve from the user points.

**Fig. S8.**
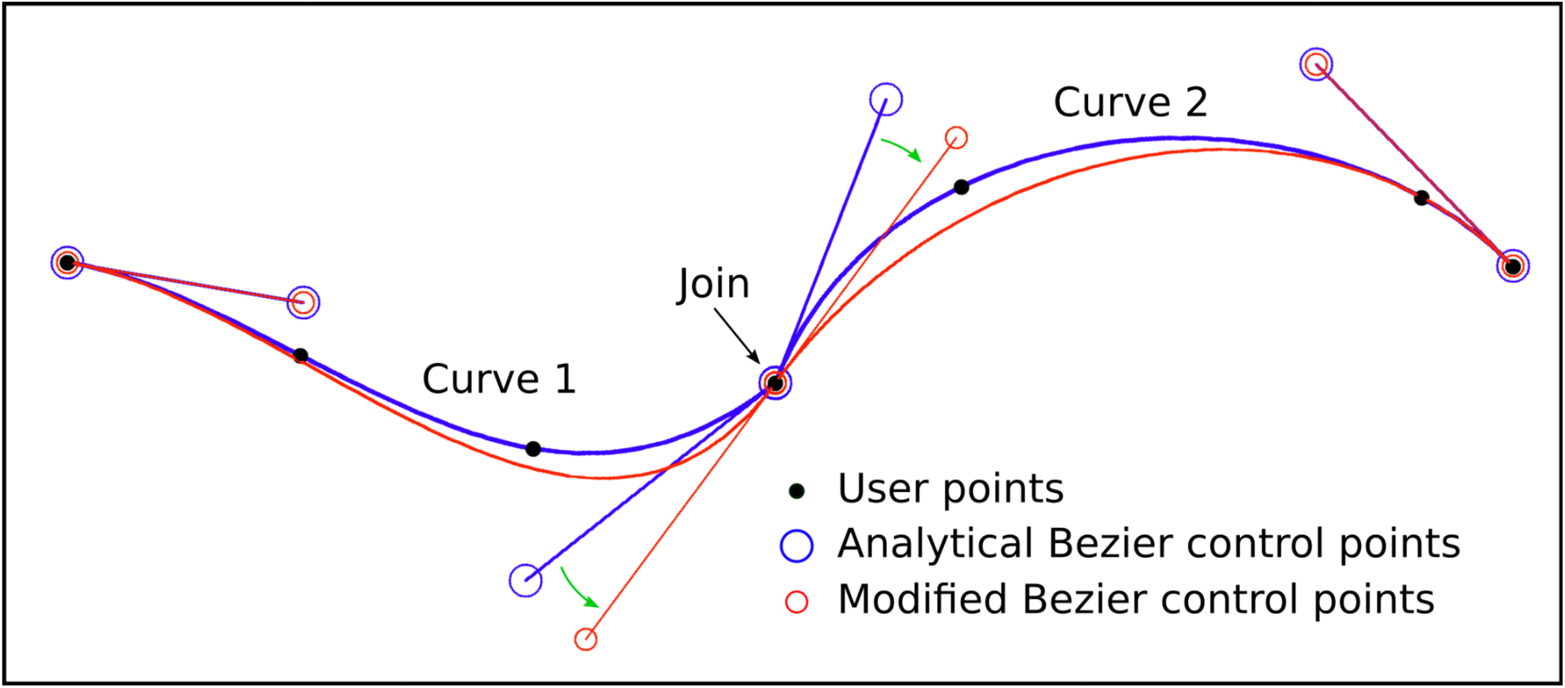
Two cubic Bezier curves are shown in blue, fit to the black, user-defined points. The blue circles are the analytically determined Bezier control points that provide the best fit to the user defined points. To eliminate the discontinuity in the gradient at the join, the two closest control points are rotated by equal and opposite angles about the join (green arrows) until they and the join lie on a straight line. The resulting, modified Bezier curve is shown in red. The modified curve no longer passes through all of the user supplied control points, but it has a smooth gradient throughout.

### Sample Boxes

Once the curves have been defined, the Stalefish visualization tools can be used to display the sample boxes. N sample boxes are defined by drawing N+1 equally spaced normal vectors from the curve. To find N+1 equally spaced locations on the curve (which is made of 1 or more individual Bezier curves) we follow the following numerical procedure:

- Compute the distance from the first point on the curve to the end point (that is, the very final point of the final Bezier curve).
- Divide this by N to get a candidate spacing, *s*.
- Up to N times: advance a Euclidean distance *s* along the curve, recording the coordinate at each step. Bezier curves are parameterized with *t* in the range [0,1], mapping coordinates on the curve from its start to its end. The increment of *t* which will advance a coordinate a distance *s* along the curve is computed via a simple binary search. The algorithm takes account of steps that cross the join of two Bezier curves.
- Review the number of coordinates that could be fit onto the full curve for spacing *s.* If the number of coordinates is different from N+1, adjust *s* (by doubling/halving it) and repeat the previous step. Repeat this step until the number of coordinates on the curve is N+1.

The start and end of adjacent vectors provide four corners of a box (Fig S9 A). Controls are provided to allow the sample boxes to extend above or below the curve (Fig S9 C/D). Note that sample boxes may overlap, if the curve is sharp and the boxes extend a long distance (Fig S9 B). In future work it may be desirable to define sample boxes between two user-defined curves to avoid this problem.

**Fig. S9.**
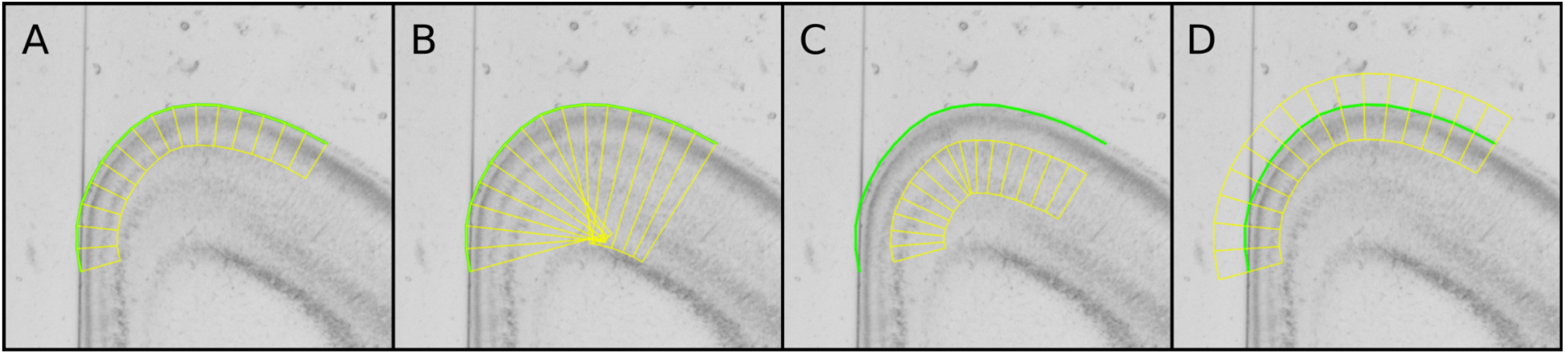
(A) 16 sample boxes along a curve comprised of three cubic Bezier sections which follows the outer edge of the isocortex. (B) When the sample boxes are over-extended, they overlap, meaning that some pixels of the image will form part of multiple sample boxes. (C) It is possible to use the same curve to sample multiple regions. This example samples a deeper region than the sample boxes in A, making use of the same curve. (D) If required, the sample boxes may be extended above as well as below the curve.

The mean signal value in the box (and its standard deviation) is computed and stored in the Stalefish project file. Optionally, the value, coordinates and in-box depth of each pixel in each sample box can be saved into the project file.

### Freehand mode

Freehand mode allows for the encircling of a region on each brain slice so that the signal encoded in pixels within the region can be saved into the HDF5 project file.

### Signal recovery

The *Stalefish* technique assumes that there is some monotonic relationship between the value of a pixel in the brain slice image and a variable of interest (*Id2/RZRB* gene expression in the current study). The value of a pixel may be a simple luminance if the slice images are greyscale, or it may be that color information needs to be accounted for, such as in the Allen Developing Mouse Brain Atlas or in certain recent multiple-ISH staining techniques. We have implemented both a luminance/greyscale color mapping and a color mapping which can be used with Allen ISH images; the choice of color mapping is selected with an entry in the project’s JSON configuration file. The Allen color mapping is described in more detail in the section ‘Allen mouse brain maps’.

The luminance-based mapping is a straightforward mapping of the 8-bit value of any of the color channels in the image file (any image format supported by OpenCV can be used including TIFF and PNG). The stains used here are darker where there are more mRNA molecules coding for the protein of interest, thus lower pixel luminance values correlate with higher signals. The simplest possible mapping would be to assign to the pixel value 0 the signal 1.0 and to the maximum pixel value 255 the signal 0. This would work well if the image capturing process guaranteed uniform illumination of the sample. We found that samples illuminated with a Zeiss KL1500 LCD light source and captured using a Zeiss AxioCam camera mounted to a Zeiss Stereo Discovery V12 microscope in our lab generated slight variations in luminance across the sample, which were significant enough to upset the signal extraction if some sample slices were imaged in one orientation (say, medial to the left and lateral to the right of the image) but others were imaged in the opposite orientation (lateral-medial) and then inverted in the software to match the medial-lateral slices. In these mirrored slices, the systematic overall illumination gradient was reversed making it difficult to compare the signal in adjacent slices. To counter for such inhomogeneities in the illumination, we adopted a post-processing approach. We make a copy of the image, blur it with a very wide Gaussian kernel, then subtract this from the image leaving the signal, *p*_s_ according to

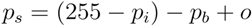

where *p*_i_ is the image pixel’s 8-bit greyscale value, *p*_b_ is the pixel value from the Gaussian blurred image and o is a constant parameter (bg_blur_subtraction_offset) chosen to keep *p*_s_ in the range [0,255]. Separate windows in the Stalefish visualization can show the blurred image and the image signal. An example is shown in Fig S10.

**Fig. S10.**
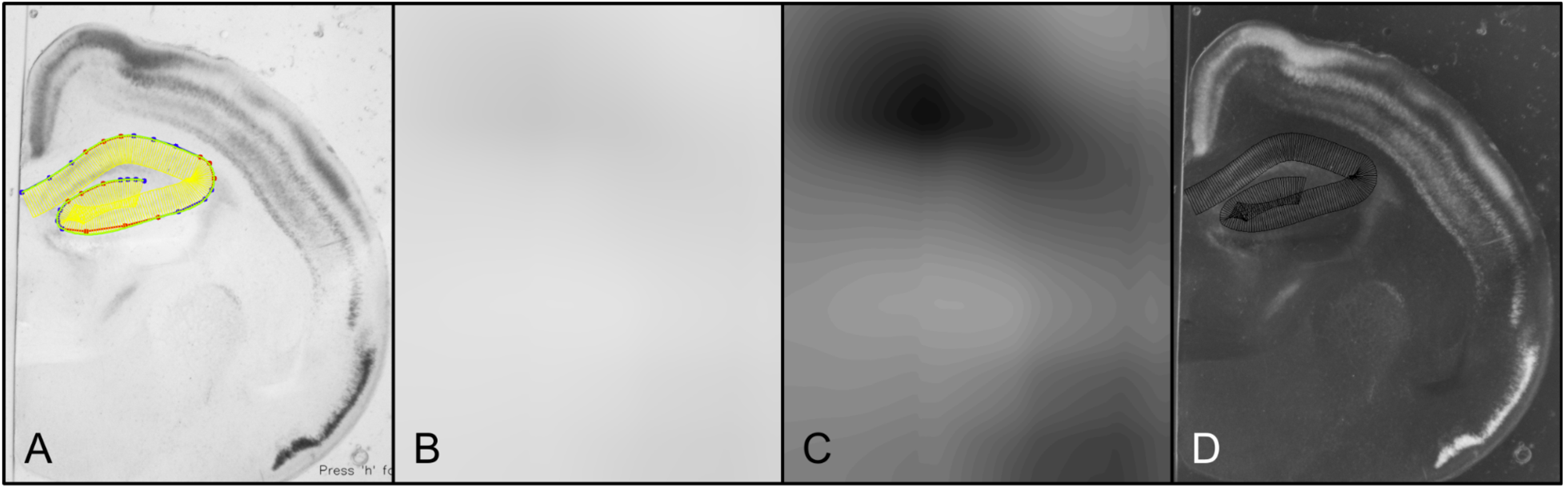
(A) The main image window shows the original Id2 ISH image of slice 26 of Vole 65_7E. This is the image upon which curves, landmarks and freehand loops are drawn. (B) Gaussian blur of A with kernel width set to 1/6 of the width of the original image. Little structure is apparent because the illumination inhomogeneities in this image are small. (C) Enhanced contrast version of B indicates that subtracting the blurred image will have a small effect on true signals as well as countering any systematic illumination inhomogeneities. (D) The signal window. Signal is drawn in greyscale with higher signals towards white and so this looks like the photographic negative of the original image. The signal window can be viewed with the ‘e’ key in Stalefish; the blurred image with the ‘r’ key. Note that the sample boxes are shown on the signal window using thin black lines.

### Landmarks

Landmarks are coordinates defined on the brain slice image matching either anatomical features or researcher-added alignment marks. We distinguish between landmarks which are expected to be found on every slice and those which are present on only one or a few slices. Landmarks present on every slice are used for slice alignment or for tracking structures when the 3D reconstruction has been made. Landmarks are added using either *Stalefish’*s ‘Landmark’ mode or in ‘Circlemark’ mode.

**Fig. S11.**
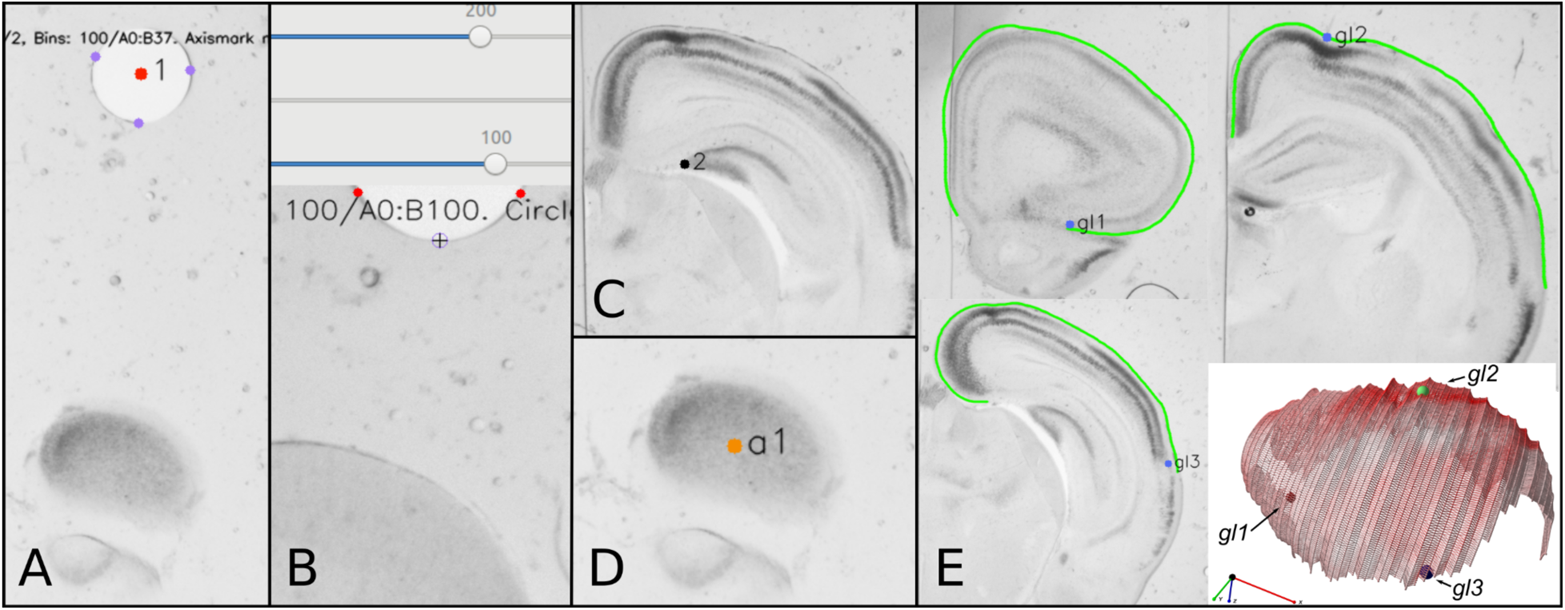
Landmarks. (A) ‘Circlemarks’: The centre of a full circle is estimated by placing three points around its circumference and finding their circumcircle. This defines a landmark, which in this slide is numbered ‘1’ as it is the first one. it is red because there is not a corresponding landmark ‘1’ on every one of the other slices in the set. (B) It is possible to estimate the circle centre even if only a part of the needle-created circle is visible in the frame. (C) A regular landmark is defined by the user as a point. This landmark is marking the dentate gyrus in the hippocampus. (D) Axismarks mark the ends of the brain axis. They are marked by orange dots. (E) Three slices from a set on which are marked 3 global landmarks. The green line of the user-defined curves are shown; note that the global landmarks are defined by anatomical features but lie close to the curve in each case. The three dimensional render of the brain shows the three landmarks as spheres. These are the three landmarks which are used to make linear transformations of the digitally unwrapped map.

### Globalmarks

’Globalmarks’ are landmarks which are used for linear transforms. Globalmarks are stored in a data structure in the HDF5 file in the order in which they were added to the project.

### Axismarks

’Axismarks’ are landmarks which define a brain axis. A defined axis which passed through a brain surface is important for the digital unwrapping of the surface. The brain axis may not be aligned with any of the coordinate axes and even if it is, the user must supply a piece of information to declare which this would be as the brain may have been coronally or sagitally sliced (our convention is to say that the brain slices lie in the y-z plan and are stacked along the x axis). The user can add two coordinates to a brain slice set using ‘Axismark’ mode to define the endpoints of a brain axis. The use of the brain axis is described in the section ‘Digital unwrapping’, below.

### Landmark Alignment

Once curves, or freehand regions have been drawn on all slices, we want to align adjacent brain images so that the aligned curves will form a three dimensional surface. The most reliable way to achieve alignment is to form visible markers in each slice preparation. As previously described, we used a needle to form circular marks on each slice. These circular marks were used for a ‘landmark alignment’ process proceeding as follows: The user marks three points on each circular landmark. The best estimate of the alignment landmark is given by the center of the circumcircle passing through the three marked points. A two-dimensional coordinate offset is applied to each slice to place the alignment landmarks in a line in 3D space that is parallel with the x-axis to form an ‘alignment axis’. Then, starting with the second slice image, each slice is rotated about the alignment axis so that the points on the curve are as close as possible to the points on the curve in the previous slice. This is determined by minimizing the sum of squared distances between N equally spaced locations on the curve on slice *i* and the corresponding N locations on the curve of slice *i*-1. We call this alignment technique ‘landmark alignment’. It is based on the assumption that the anatomist has marked curves which correspond to the same anatomical structure on each brain slice.

We note that the best alignment accuracy using a landmark based alignment method would be to form *two* needle-formed alignment marks in the fixing medium and then perform an affine transformation of each slice image to align both alignment marks. This is left as a future enhancement to be developed in the software.

### Auto Alignment

As an alternative to landmark alignment, if an existing slice set is to be analysed which does not have the necessary alignment marks, we developed an ‘auto-alignment’ algorithm. This uses only the two dimensional sample curves on each slice. For each slice, *i*, a translation, **r** and rotation, ϕ, are found which will position the curve points optimally with respect to the previous slice and a single ‘target’ slice. We used a Nelder-Mead optimization process (*30*), which finds a minimum for the following cost function:

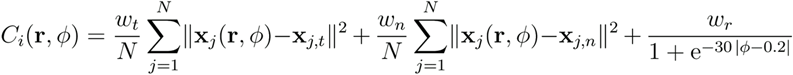

where the first term computes the sum of squared distances between *N* transformed candidate points, ***x***_j_ (which are evenly distributed points on the curve of slice *i* that have had the translation ***r*** and rotation ϕ applied) and N candidate points ***x***_j,t_ which are evenly distributed points on the target (middle) curve in the slice set; the second term computes the sum of squared distances between ***x***_j_ and N neighboring points, ***x***_j,n_ on slice *i*-1. The first term ensures that the slice positions do not ‘drift’ by penalizing large translations away from the centroid of the target slice. The second term ensures that each slice is closely aligned to its neighbor and the third term penalizes large rotations of any curve; it is a sigmoid curve whose parameters were set by hand to penalize rotations greater than about 0.2 radians, without affecting small rotations. *w*_t_, *w*_n_ and *w*_r_ are weights with the values 0.01, 1 and 0.1, respectively.

### Software dependencies

Stalefish was developed using the image processing library OpenCV (*31*) together with Bezier curve processing features and other supporting code from morphologica (https://github.com/ABRG-Models/morphologica). OpenGL-based visualization in the tool sfview is also provided by morphologica.

### Data Analysis

This section describes how the data generated from a Stalefish project - essentially a set of mean expression values with spatial coordinates - can be rendered as a three dimensional image or converted into a two dimensional expression map. All data for a project is written into a single project file, whose format we discuss first.

### Project file format

Stalefish writes data in Hierarchical Data Format, version 5 (HDF5), a global standard file format. HDF5 files can be read with a multitude of software tools and code libraries, including Python, R, MATLAB, GNU Octave, C and C++. The HDF5 project file is named to match the JSON configuration file from which the project was created. Thus, if the JSON file is called Mouse_DS4.json, then the resulting HDF5 project file will be named Mouse_DS4.h5. The HDF5 format is standard, but the choice of variable containers in an HDF5 file is application specific.

Data variable names in an HDF5 file look very much like folder paths on a computer filesystem and we refer to HDF5 variables as being contained in ‘folders’. The data in a Stalefish project is divided into numbered folders; one for each slice frame; the first frame is contained in /Frame001, the second in /Frame002; etc. Each Frame folder contains a number of sub-folders containing location information, and a sub-folder which contains the extracted signal information. There is a full description of the Stalefish HDF5 format at https://github.com/ABRG-Models/Stalefish/tree/master/reading with example Python and GNU Octave code for reading (and plotting) the data available from the same location.

### 3D Brain

The stalefish program allows the annotation of a set of brain slices, and saves information about the aligned data into an HDF5 file. To view and manipulate 3D renderings of the data in the HDF5 file, we wrote a simple viewer application called sfview, controlled by command line arguments. sfview can be used to visually inspect and verify the quality of the alignment of a set of slices and also to transform a set of digitally unwrapped surfaces onto a single individual example, writing out the transformed ‘digital unwraps’ into separate HDF5 files.

To render a gene expression surface, we must decide how to plot the mean signal value for each sample box. We have used two methods. In each method, we use the sample box vertices that lie *on* the curve. The first method uses these vertices to define a series of ‘ribbons’, one for each brain slice. This view, shown in Fig S12 A & B, is useful for analyzing how well the chosen alignment algorithm has arranged the slices and how the curve shape progresses across the sample. The second method takes the on-curve sample box vertices and uses these to define a triangular mesh (Fig S12 C). This results in a smoother expression surface (Fig S12 D).

**Fig. S12.**
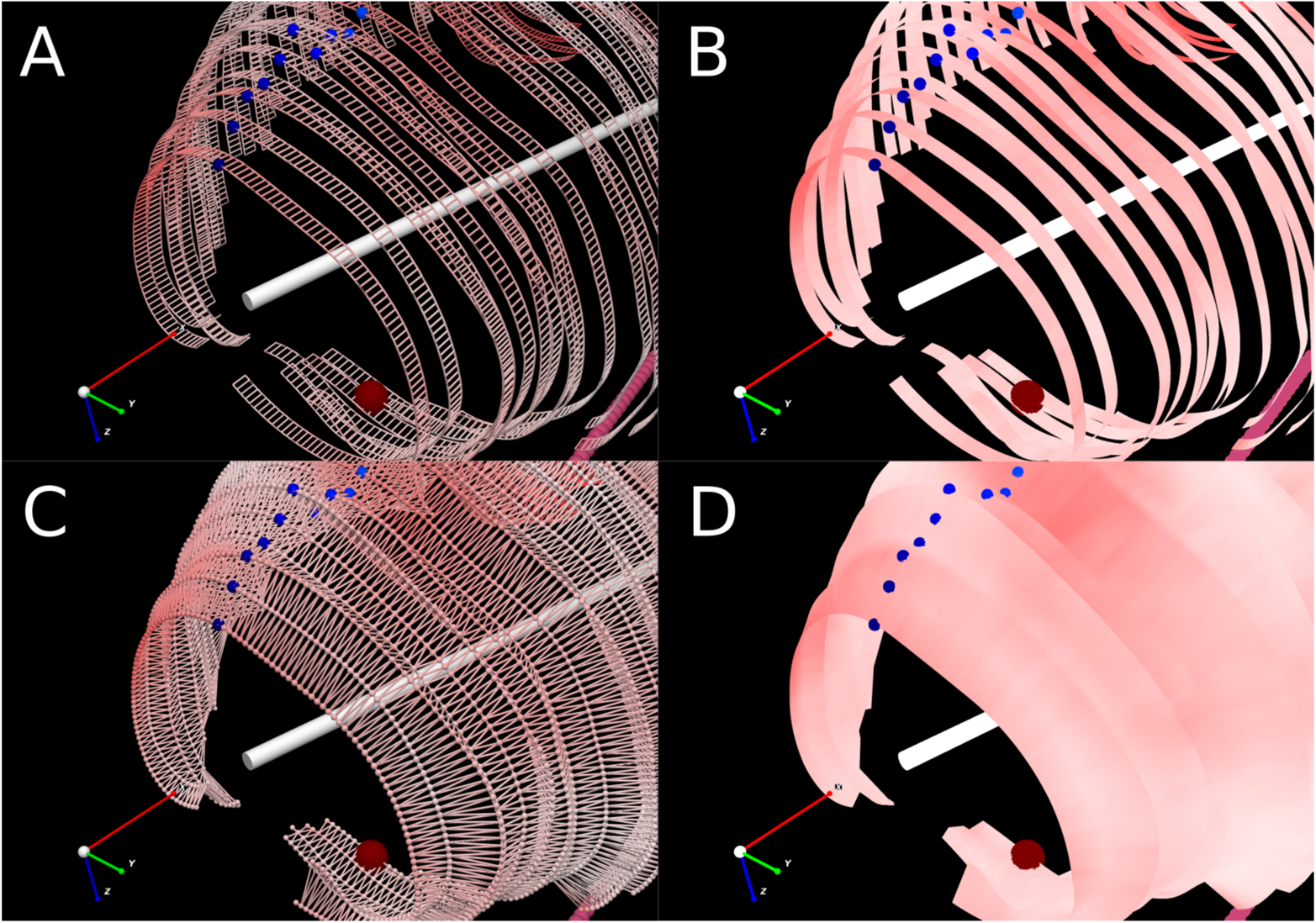
Two ways to render the mean expression for the samples boxes into a three dimensional view. In each view the user-defined brain axis is shown as a white bar, the digital unwrapping ‘zero marks’ are shown as a row of small blue spheres and a landmark is displayed as a larger, burgundy sphere. The straight row of alignment landmarks is visible in pink at the bottom right of each panel. (A) Use two sample box vertices (each with a mean expression value) that lie on the curve, and extend along the x axis by the slice thickness to define two more vertices, forming a rectangular region of expression. The edges of each rectangle so defined are shown here to illustrate. The expression signal is shown using the color red, with the highest signal given by the most saturated red regions, but note that here, a shader that provides a diffuse lighting effect has been used and this distorts the expression colors slightly. (B) The same ‘ribbon’ view of the slice data, where color is defined at each vertex, but varied linearly across each rectangle (a task performed automatically by the OpenGL shader). (C) To produce a smoother surface, we use the sample box vertices on each curve to define a triangular mesh.

Here, the mesh is illustrated with lines and spheres. (D) The smoothed version of C, with OpenGL performing color interpolation between the vertices as in B.

### Digital unwrapping

Digital unwrapping is the process of straightening out a curved, three dimensional surface into a two dimensional map. The process begins with a set of aligned curves, as in Fig S13 A. The user provides axismarks that define a brain axis (white bar). An unwrapping axis of ‘zero marks’ is defined on the surface, by rotating a user-defined angle about the x-axis (centered on the brain axis), then locating the most distal point on each curve at this angle (blue/rainbow spheres in Fig S13 A). Each expression ‘ribbon’ is now straightened out, holding it fixed at its zero mark (Fig S13 B). In Fig S13 C, the straightened ribbons have been inverted and a further rotation shown in Fig S13 D shows that the data begin to resemble the two dimensional map in Fig S13 E, in which the zero marks have been arranged to lie on a straight line, which means that there are now no gaps between the ribbons. Note that the quadrilaterals which make up the ‘pixels’ in Fig S13 E are not of even size; those in the shorter ribbons are smaller than those in the longer ribbons (because in this example, there are the same number of sample boxes on each curve/ribbon).

Furthermore, although this particular map has not been transformed; it is possible that a transformation may be applied to the map in Fig S13 E, transforming rectangular pixels into general quadrilaterals prior to resampling. The final step is to resample the image in Fig S13 E to produce an image consisting of square pixels, as shown in Fig S13 F.

**Fig. S13.**
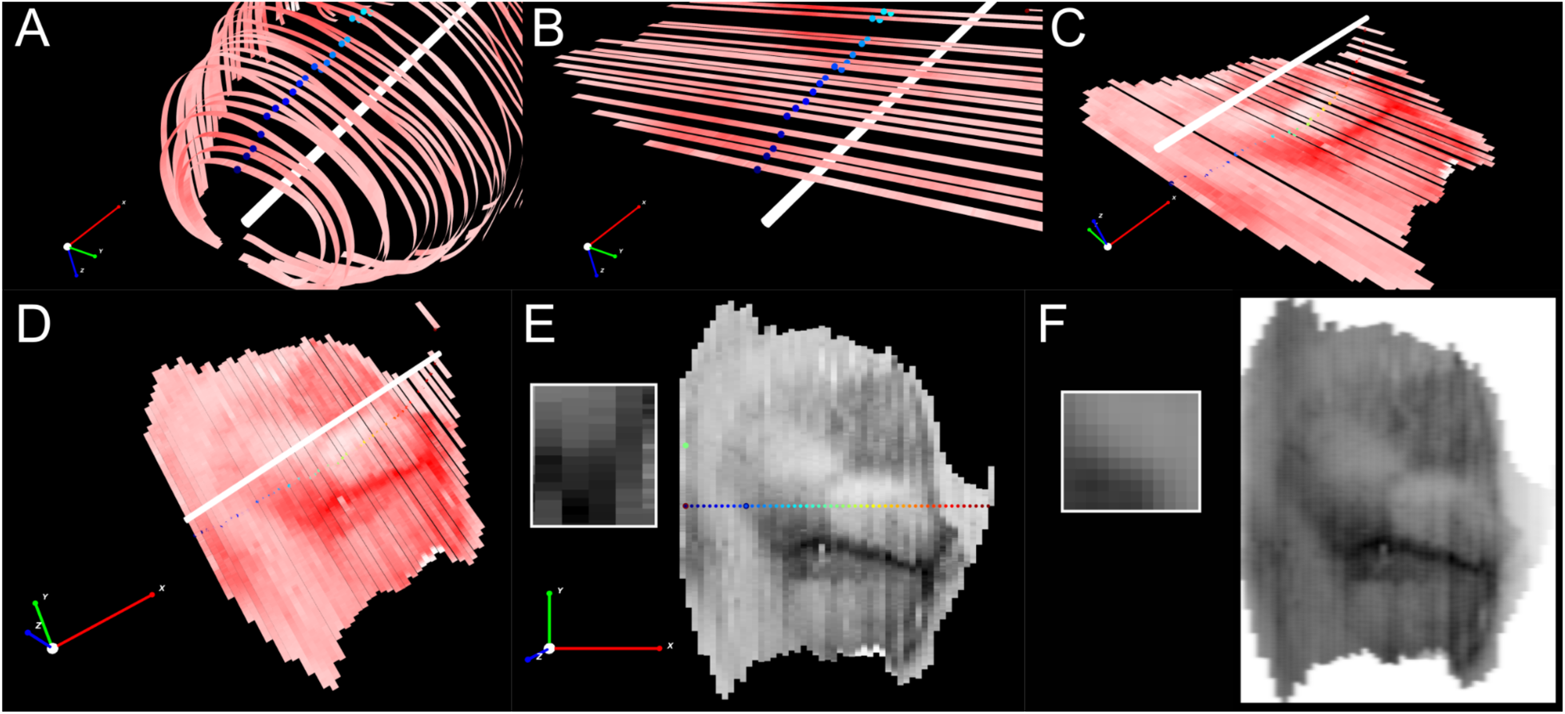
The digital unwrapping process. (A) To illustrate the process, we start with a Vole brain with Id2 expression shown as ‘ribbons’ which follow the curves defined in Stalefish. The brain axis is visible as a white bar and at a fixed angle about the x axis, a series of ‘zero marks’ are shown on the ribbons as rainbow colored spheres. (B) The ribbons are straightened out using the zero marks as fixed points. The 3D view in A and B is identical; the brain axis and zero marks are unmoved. (C) The view is zoomed out and inverted with respect to panel B (compare the xyz coordinate arrows) (D) Further rotation of the 3D view. The gene expression pattern is now visible. (E) The unwrapped ribbons are now aligned by taking the zero marks and arranging them along a straight line. Note that this image still consists of quadrilaterals of varying size (inset). There are as many quadrilaterals in the short end ribbons as in the long central ribbons. (E) The image is resampled using a sum of Gaussians method to produce the final image, whose pixels are now square (inset).

The resample algorithm finds a signal value, p*_k_*, for each square pixel in a resampled grid (fig. S13 F). *p*_k_ is a sum determined from the contributions of *M* quadrilaterals indexed by *j*, in each of *N* ribbons according to a 2D elliptical Gaussian distribution centered on each quadrilateral. The parameters of each elliptical Gaussian are determined by the shape of the quadrilaterals. This can be expressed as

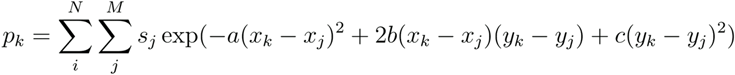

where *s*_j_ is the signal of quadrilateral *j*, (*x*_k_, *y*_k_) are the coordinates of the square pixel; (*x*_j_, *y*_j_) are the coordinates of quadrilateral and a, b and c are given by

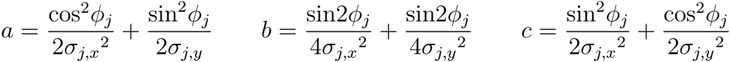

where σ_j,x_ and σ_j,y_ are the parameters of an ellipse rotated by the angle ϕ_j_. We used 3 corner coordinates of the quadrilateral (**c**_1_, **c**_2_ and **c**_3_) to determine these parameters. Suitably chosen, these give two basis vectors, ***x***’ = **c**_3_ - **c**_2_ and ***y***’ = **c**_1_ - **c**_2_ for the quad which define the major and minor axes of the ellipse:

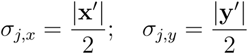

The rotation of the ellipse, ϕ_j_, is defined as the angle which ***x***’ makes with respect to the x axis, i.e.

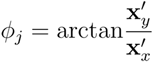

### Protocol for processing images to generate 3D and 2D surface expression maps

- Create a text file with a .json suffix and populate it with the mandatory elements given in Table 1 and with reference to the example in Fig S14.
- Launch Stalefish with the path to the .json file as a single argument. It will load the images and present the first one to the user in two windows, one a ‘working’ window and a second which displays the mRNA signal to the user (after subtracting the blurred background).
- Cycle the input mode to ‘Circlemark’ mode (see Table 2 for a list of Stalefish functions). This allows the location of the alignment needle mark to be set for each slice. Place three marks around the circular boundary of the needle hole allowing the program to mark the center of the hole. Repeat for each slice in the set.
- Cycle the input mode to ‘Axismark’ mode. This is used to define a central axis through the brain samples. Mark exactly two axis marks in the entire slice set.
- Cycle to ‘Curve’ mode. Using the mouse, place 3 or 4 points to define a part of the curve on the slice. Press space to commit a curve portion; it will turn red or blue. Cancel points and replace them as necessary until the curve portion follows the anatomical structure satisfactorily. Define 3 or 4 more points along the curve and press space to commit a new curve portion. Continue until a full curve has been defined for the structure of interest. Repeat for all brain slices.
- Cycle to ‘Global landmark’ mode. Define exactly three global anatomic landmarks across all of the slices in the brain. Each landmark should ideally be relatively close to the curve.
- Use the save function to write the data to an HDF5 file (the structure of the data content in this file is described separately). Exit Stalefish.
- If analyzing a single brain, open the HDF5 file using the sfview program to verify that the slice alignment and 2D map unwrapping was successful. Use the −m1 argument to view the 2D unwrapped map. For example, if the json file was named brain1.json, the HDF5 file will have been named brain1.h5 and the correct sfview command would be ‘./build/src/sfview brain1.h5 - m1’
- If analyzing several brains, then follow steps 1 to 7 to define curves and global landmarks on each brain. The brain maps can be transformed onto the same coordinate axes using sfview’s-T argument, which computes transformations based on the 3 global landmark coordinates provided on each brain. For three brains, an example sfview command is:

*./build/src/sfview brain1.h5 brain2.h5 brain3.h5 -m1 -T*

**Table 1:**
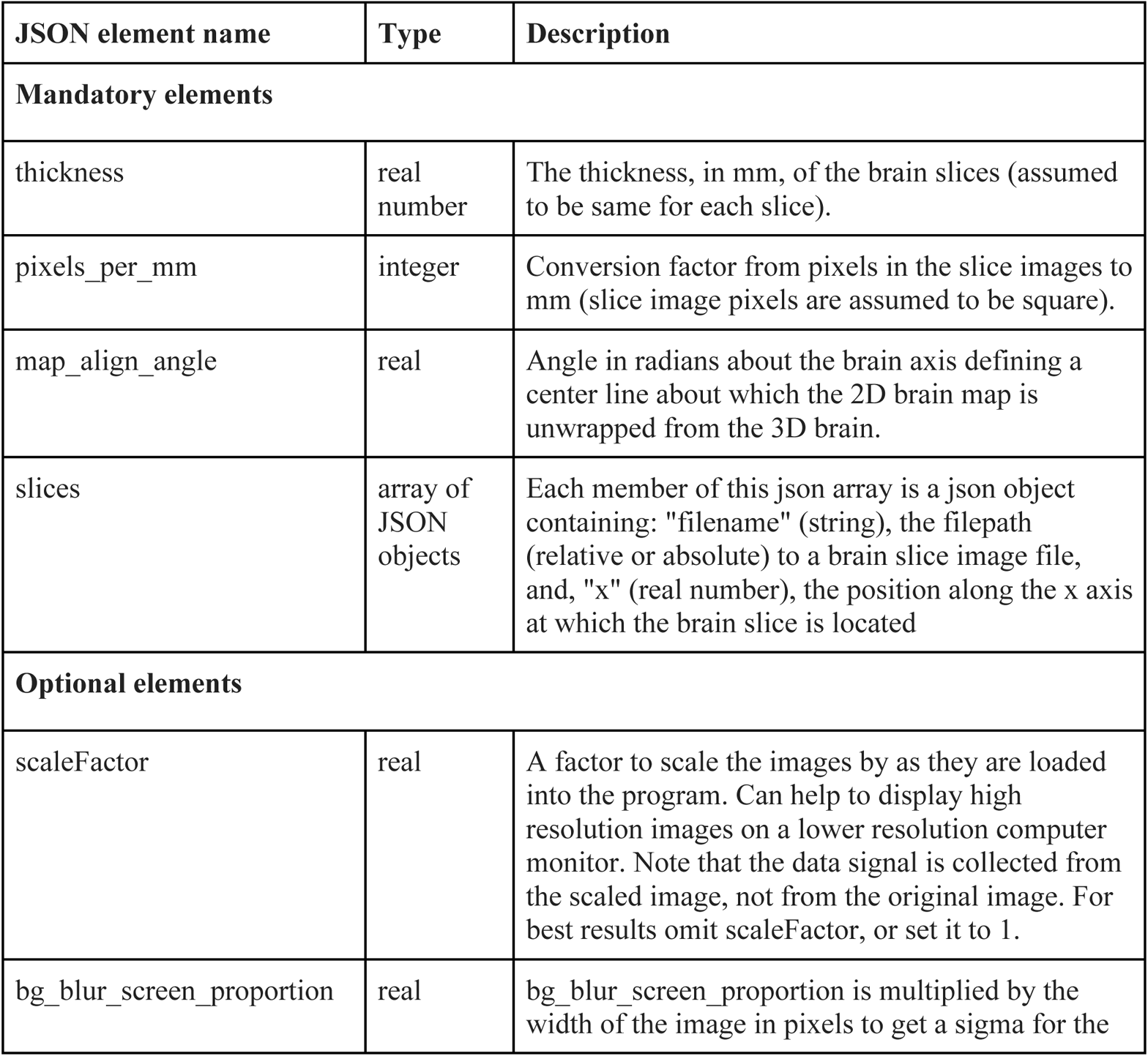
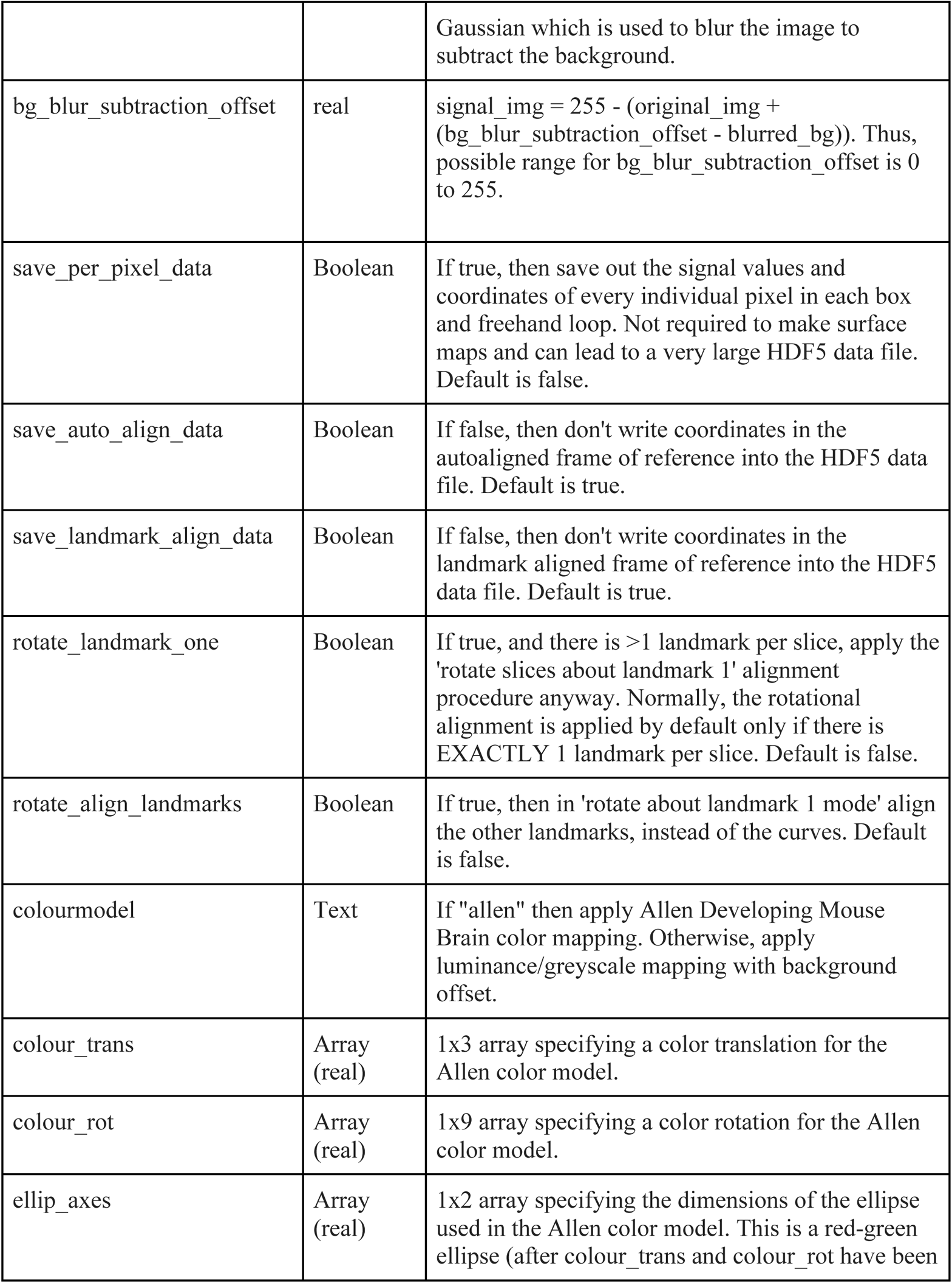
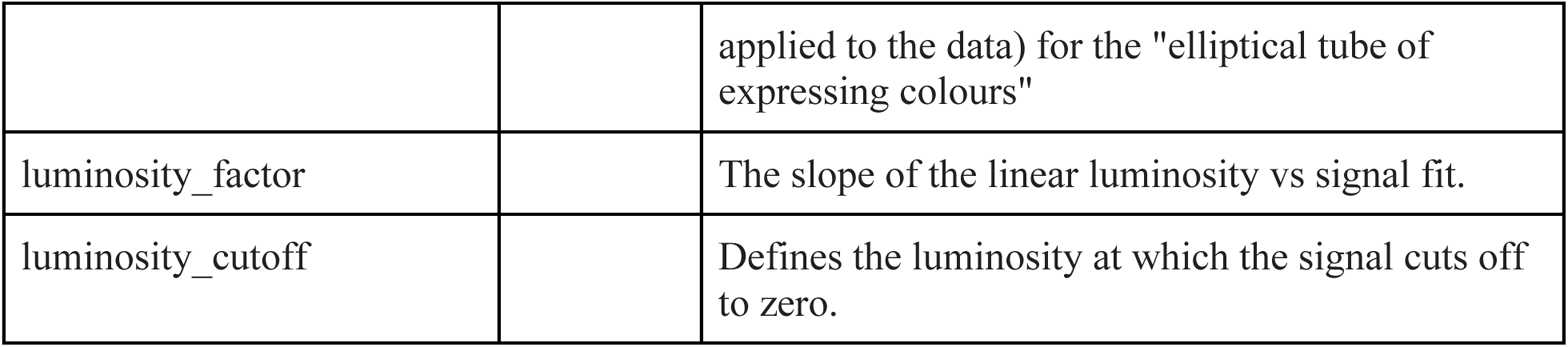
mandatory and optional parameters which should be written into a Stalefish project’s JSON configuration file.

**Table 2:** Stalefish functions

| Function | Key | Description |
| --- | --- | --- |
| Box A | user interface slider | 'Sample box position A'. Change the position of the start of the sample boxes |
| Box B | slider | 'Sample box position B'. Change the position of the end of the sample boxes |
| Num bins | slider | Change the number of sampling bins on the curve (range: 2-200) |
| Toggle Bezier control points | 1 | Toggles the visibility of the Bezier control points for the curves |
| Toggle user points | 2 | Hides/shows the user-supplied control points |
| Toggle fit line | 3 | Hides/shows the green line of best fit to the user-supplied control points |
| Toggle the bins | 4 | Hides/shows the yellow sample bins |
| Move to next curve | Space | Commits the green 'pending' user points to be a Bezier curve; new user points will become part of the next curve. |
| Cancel | c | Cancel the last point or freehand region, depending on context. If in *Curve mode* it cancels the last user-supplied curve point; if in *Globalmark mode* then the last global landmark is cancelled. |
| Delete all curves | C | Clear all user-supplied points from the current project |
| Update fit | f | Recompute the Bezier fit on the current frame |
| Update fit (all frames) | F | Recompute the Bezier fit on the all frames |
| Copy bin params | B | Copy the sample bin params to all other frames |
| Write file/Save | w | Write the project to an HDF5 file |
| Cycle mode | o | Cycles the input mode between: 1) Curve mode; 2) Freehand mode; 3) Landmark mode; 4) Globalmark mode; 5) Circlemark mode; 6) Axismark mode. |
| Curve mode: start/end | s | When in curve mode, this switches the input to add (or cancel) user points to either the start or the end of the curve. Adding at the end is default. |
| Export user points to tmp | k | Export data to /tmp/landmarks.h5, /tmp/curves.h5 and /tmp/freehand.h5 |
| Export landmarks etc to tmp | p | Export mode-specific data to /tmp |
| Import landmarks | l | Import landmarks from /tmp/landmarks.h5 |
| Import curve points | i | Import curve points from /tmp/curves.h5 |
| Import freehand | j | Import freehand loops from /tmp/freehand.h5 |
| Next frame | n | Move to the next frame in the brain slice set |
| Back to previous frame | b | Move to the previous frame in the set |
| Mirror frame | m | Mirror the image (left-right) |
| Toggle blur window | r | Toggle the window that shows the Gaussian-blurred background |
| Toggle signal window | e | Toggle the window that shows the final signal |
| Help | h | Displays a summary of the key functions on screen |
| Exit | x | Exit the program |

In this case, brain2 and brain3 would be linearly transformed to match brain1 (which would not be transformed). To transform to match brain2, brain2.h5 would be given as the first argument. sfview will write out the transformed and resampled data into separate .h5 files with a naming scheme showing which 2D brain map is stored and from which brain its transformation was computed (’Transformation From’). The example above would result in these files:

brain1.TF.brain1.h5

brain2.TF.brain1.h5

brain3.TF.brain1.h5

**Fig S14.**
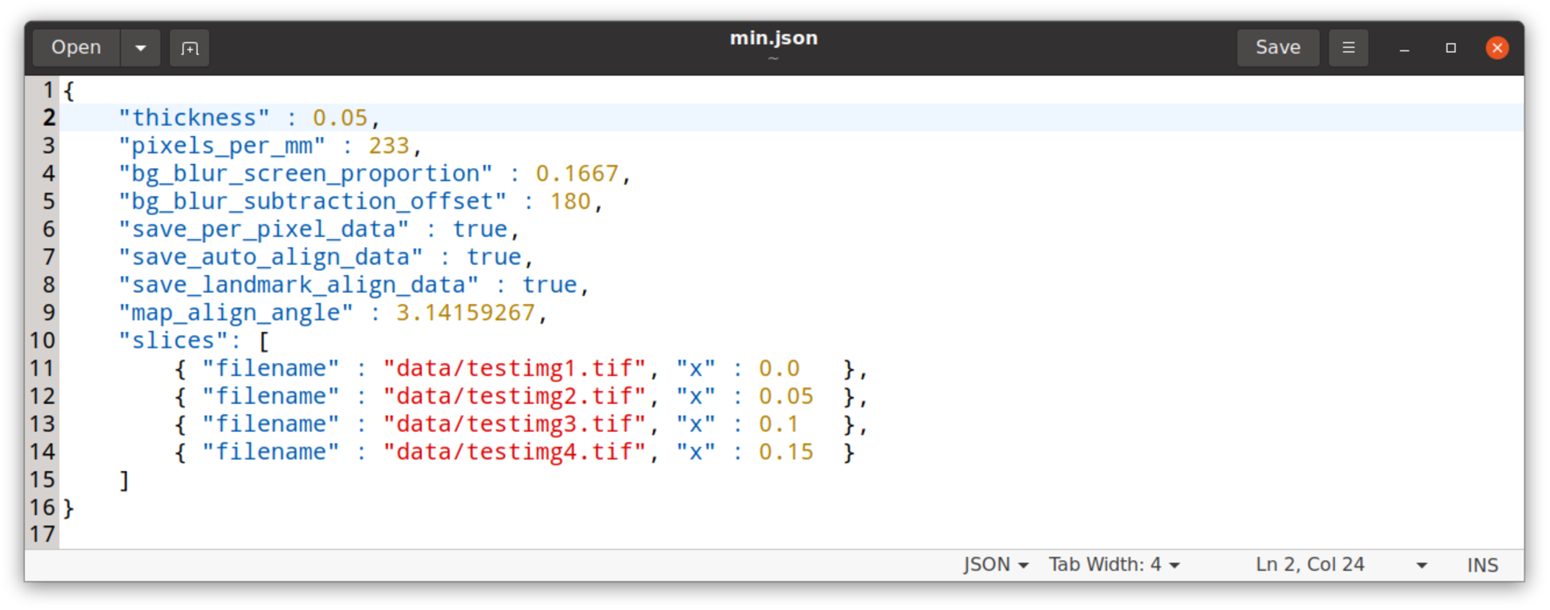
An example Stalefish JSON configuration file. This example contains the mandatory elements, plus a few of the optional elements. The project contains four slice images.

**Table 3:** sfview functions

| Function | Key | Description |
| --- | --- | --- |
| Help | h | Output a helpful key summary to stdout |
| Exit | x | Exit sfview |
| Scene lock | l | Toggle the scene lock to prevent mouse movements from changing the view |
| Mouse rotate mode | t | Toggle the axes about which the scene is rotated for mouse movements |
| Coord arrows | c | Show/hide the small coordinate arrows |
| Snapshot | s | Take a snapshot, which will be saved as picture.png |
| Reset view | a | Reset to the default viewpoint |
| Reduce field of view | o | Reduce the field of view of the virtual camera |
| Increase field of view | p | Increase the field of view of the camera |
| Save view | z | Saves the current view location to a temporary file, which will be restored on future runs of the program |
| Reduce the near cutoff plane | u | Reduce the plane beyond which objects are rendered |
| Increase near cutoff plane | i |  |
| Select model | 0-9 | When sfview is showing multiple expression surfaces, they can be individually selected with the number keys |
| Decrease opacity | Left | Decrease the alpha channel value for the selected model, making it more transparent |
| Increase opacity | Right | Increase the alpha channel value for the selected model, making it more opaque |
| Toggle landmarks | f | Show/hide the landmark locations |
| Toggle zero angle marks | g | Show/hide the zero angle marks which are used to digitally unwrap a 3D surface into a 2D map |
| Toggle brain axis | d | Show/hide the user-defined brain axis |
| Toggle 2D map | j | Show/hide the unwrapped 2D map |
| Toggle 3D map | k | Show hide the 3D expression surface |

### Future development of *Stalefish*

*Stalefish* is a simple tool, implementing the well-defined and limited set of algorithms described in this work. We followed the design philosophy of doing a simple thing and doing it well. As such, we hope that significant future development of the software will not be necessary.

However, some improvement may be desirable in the algorithm which joins separate Bezier curves together, with a more sophisticated optimization applied to the way that the algorithm adjusts the adjoining curves’ control points.

The data generated by *Stalefish* is intended to be rendered in any tool of the researcher’s choice. One possibility would be to use the recently developed BrainRender (*33*), though we have not yet investigated any changes that may be necessary to make this work.

An aspect of the *technique* that warrants future development is the definition of anatomical landmarks and the way that three dimensional surfaces are digitally unwrapped. We have one feature in particular in mind; ‘manual unwrapping’. Here, rather than using the angle about the brain axis to define the ‘zero marks’ about which the surface is unwrapped, it might be possible to use manually placed landmarks on each brain slice. This might work for the Hippocampal data in Fig. S4 for which the dentate expression on each slice could be used to define the unwrap ‘zero-mark’.

